# The telomeric Cdc13-Stn1-Ten1 complex regulates RNA polymerase II transcription

**DOI:** 10.1101/493296

**Authors:** Olga Calvo, Nathalie Grandin, Antonio Jordán-Pla, Esperanza Miñambres, Noelia González-Polo, José E. Pérez-Ortín, Michel Charbonneau

## Abstract

Specialized telomeric proteins have an essential role in maintaining genome stability through chromosome end protection and telomere length regulation. In the yeast *Saccharomyces cerevisiae*, the evolutionary conserved CST complex, composed of the Cdc13, Stn1 and Ten1 proteins, largely contributes to these functions. Here, we report the existence of genetic interactions between *TEN1* and several genes coding for transcription regulators. Molecular assays confirmed this novel function of Ten1 and further established that it regulates the occupancies of RNA polymerase II and the Spt5 elongation factor within transcribed genes. Since Ten1, but also Cdc13 and Stn1, were found to physically associate with Spt5, we propose that Spt5 represents the target of CST in transcription regulation. Moreover, CST physically associates with Hmo1, previously shown to mediate the architecture of S phase-transcribed genes. The fact that, genome-wide, the promoters of genes down-regulated in the *ten1*-*31* mutant are prefentially bound by Hmo1, leads us to propose a potential role for CST in synchronizing transcription with replication fork progression following head-on collisions.

## Introduction

Telomeres consist of an elaborate, high-order assembly of specific TG-rich repetitive DNA sequences and proteins that cooperatively provide protection against chromosome degradation. A number of telomeric proteins have been identified and, together, they act to “cap” the telomere and “hide” it from cellular DNA repair, including recombination (de Lange 2009). If left unprotected, telomeres are recognized by the cell as DNA double-strand breaks, leading to recombination, chromosome fusions and broken and rearranged chromosomes. Telomeric DNA is replicated by a specialized reverse transcriptase enzyme, telomerase. In addition, telomeres recruit specialized proteins to prevent telomere degradation and, hence, chromosome erosion, and regulate telomere length, including through the recruitment of telomerase at telomere ends. In vertebrates, telomere protection is provided mainly by shelterin, a complex of six telomeric proteins, TRF1, TRF2, POT1, TIN2, TPP1 and RAP1 (Palm and de Lange 2008; Martinez and Blasco 2011). A similar complex exists in the fission yeast *Schizosaccharomyces pombe* (Miyoshi et al. 2008), while in the budding yeast *Saccharomyces cerevisiae* a somewhat simpler telomeric complex, called CST, consisting mainly of the Cdc13, Stn1 and Ten1 proteins is present (Garvik et al. 1995; Grandin et al. 1997, 2001). On the other hand, recently, orthologs of *S*. *cerevisiae* CST have been found in humans and mouse, as well as in *S*. *pombe* and the plant *Arabidopsis thaliana* (Martin et al. 2007; Miyake et al. 2009; Surovtseva et al. 2009). Recently, hCST was found to associate with shieldin at damaged telomeres to regulate, in association with Polα, the fill-in of the resected overhangs and facilitate DNA repair (Mirman et al. 2018). In yeast, Stn1 has also been implicated in the fill-in of the strand previously elongated by telomerase (Grossi et al. 2004; Lue et al. 2014). Based on the hypersensitivity of mutants of CST to DNA damaging agents and its presence at sites other than the telomeres, hCST has emerged as an important potential player in counteracting replication stress genome-wide (Miyake et al. 2009; Stewart et al. 2012; Kasbek et al. 2013).

Transcription by RNA polymerase II (RNA pol II) is achieved through different steps (preinitiation, initiation, elongation and termination), and is highly regulated by a huge number of factors, including general transcription factors, cofactors, elongation and termination factors. Over the last decade, transcription elongation has revealed to be also a crucial and strictly regulated step (Pelechano et al. 2009). Among RNA pol II regulators, Spt5/NusG is the only family of transcription factors that has been evolutionary conserved, from Bacteria to Eukarya. In Eukarya and Archea, Spt5 forms a heterodimeric complex with Spt4 (Hartzog and Fu 2013). Spt4/5 associates with genes from downstream of the transcription start site to the termination sites, with a distribution pattern similar to that of RNA pol II (Mayer et al. 2010). Accordingly, Spt4/5 associates with RNA pol II in a transcription-dependent manner (Tardiff et al. 2007). In addition, Spt4/5 links the activities of the transcription elongation complex to pre‐mRNA processing and chromatin remodeling (Liu et al. 2009; Zhou et al. 2009). Although there has been until now no functional evidence for a role of Spt5 in connecting transcription with other nuclear processes, it is nevertheless noticeable that the DNA polymerases subunits Pol1 and Pol2 were identified as Spt5-associated proteins (Lindstrom et al. 2003).

Phosphorylation of the C-terminal domain (CTD) of RNA pol II largest subunit, Rpb1, which consits of an evolutionary conserved repeated heptapeptide motif (Tyr1-Ser2-Pro3-Thr4-Ser5-Pro6-Ser7), regulates RNA pol II transcription at several levels (Eick and Geyer 2013; Heidemann et al. 2013). CTD-Ser2 and -Ser5 phosphorylation (Ser2P and Ser5P) appear to be the most frequent modifications (Suh et al. 2016, Schüller et al. 2016). Ser2P is the mark of the elongating polymerase, while Ser5P marks the initiation step. The large number of possible CTD modifications generates a “CTD code” that coordinates the recruitment of numerous factors essential for transcriptional efficiency, RNA processing and connects transcription with other nuclear processes (Buratowski 2003, 2009; Hsin and Manley 2012). In *S. cerevisiae*, four cyclin-dependent kinases, Srb10, Kin28, Ctk1, and Bur1 (Meinhart et al. 2005; Phatnani and Greenleaf 2006) and four phosphatases, Rtr1, Ssu72, Glc7, and Fcp1 (Schreieck et al. 2014; Jeronimo et al. 2013) determine CTD phosphorylation along the transcription cycle. During early elongation, Bur1 phosphorylates CTD-Ser2 and Spt5 nearby the promoters (Qiu et al. 2009; Zhou et al. 2009), while Ctk1 phosphorylates Ser2 later during elongation, its activity being required for termination and 3’-end processing (Ahn et al. 2004). Fcp1 dephosphorylates Ser2P and its activity opposes that of Ctk1 to ensure proper levels of Ser2P during elongation and RNA pol II recycling (Cho et al. 2001).

In this study, we have uncovered specific genetic interactions between *TEN1* and several genes coding for transcriptional regulators, such as *BUR1*, *FCP1*, *SPT5* and *RPB1*. We demonstrate that Ten1 physically interacts with Spt5 and regulates its association with chromatin during active transcription. Stn1 and Cdc13 were also found to exhibit physical interactions with Spt5. Moreover, genome wide data show that the *ten1-31* mutation altered RNA pol II gene occupancy, as demonstrated by ChIP-qPCR and ChIP-seq data. Additionally, we found that Ten1 physically interacts with Hmo1, previously implicated in transcription regulation, as well as in solving difficult topological contexts when transcription has to face incoming replication forks (Bermejo et al. 2009). Based on our data, we propose a working model in which CST, traveling with the replication fork, could stimulate the restart of the transcription machinery following head-on collisions with the progressing replication forks.

## Results

### *TEN1* genetically interacts with *BUR1* and *CAK1*

In contrast with *S. cerevisiae* Cdc13 and Stn1 that have been attributed major specific functions in telomere protection and length regulation, Ten1, also implicated in these pathways, has no known specific function besides being attached to Stn1. To know more about Ten1, we set out to design genetic screens, using three different *ten1* temperature-sensitive mutants, aiming at identifying mutants that aggravated the growth defects of these *ten1* mutants at 36°C. Following screening of ~ 40,000 colonies of UV-mutagenized *ten1* strains, only one, the so-called *ten1*-*33* mut. #27 double mutant, satisfied several genetic criteria (see Materials and methods). Following transformation of this double mutant with a genomic DNA library, a clone that suppressed the aggravated growth arrest at 36°C was isolated and the rescuing activity shown to be at the *SGV1*/*BUR1* locus (**Fig. 1A**; see also **Supplemental material**). *CAK1* also isolated in the same complementation experiment (**Fig. 1A**) was only acting as an extragenic suppressor (see **Supplemental material**), in agreement with the previous finding that *CAK1* is a high-copy suppressor of a *bur1* mutation (Yao and Prelich 2002). However, like *bur1*-*80*, *cak1*-*23* also exhibited synthetic growth defects with *ten1*-*31* (**Fig. 1B; Supplemental material** and **Fig. S1A**).

**Figure 1.**
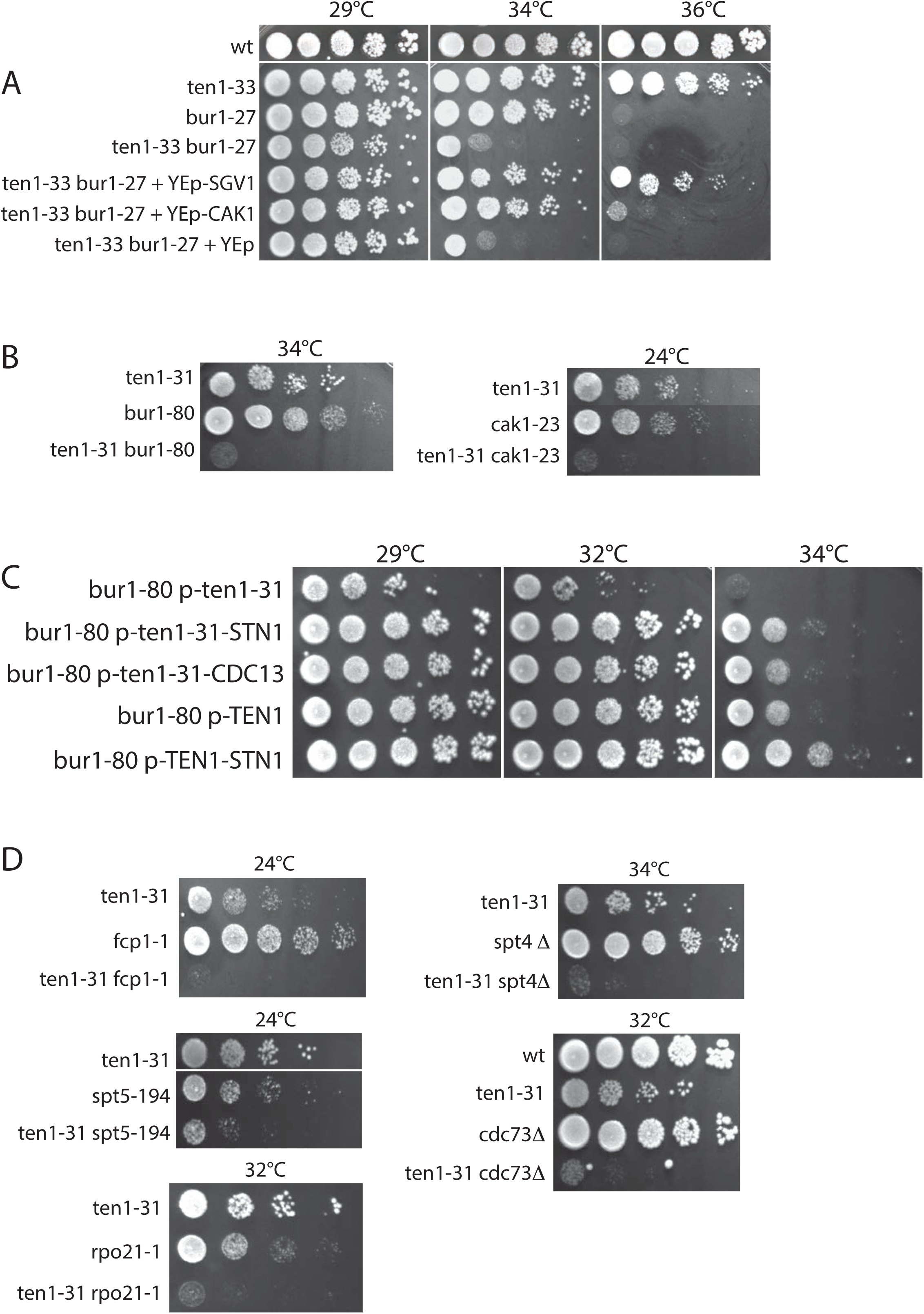
*TEN1* genetically interacts with the RNA pol II transcriptional machinery. **(*A*)** The *ten1*-*33* mutant exhibits synthetic growth defects in combination with *bur1*-*27*, as seen when comparing row 2, *ten1-33*, with row 4, *ten1*-*33 bur1*-*27*. The aggravated growth defect of the *ten1*–*33 bur1*-*27* double mutant could be complemented by overexpressing either *SGV1*/*BUR1* (row 5) or *CAK1* (row 6) from a YEp24 genomic library; in row 7, the double mutant contains vector alone as a control. Row 3 illustrates the temperature-sensitivity defect at 36°C of *bur1*-*27* alone. **(*B*)** *BUR1* and *CAK1* genetically interact with *TEN1*. Only the most relevant growth temperatures for each mutant, *bur1*-*80* or *cak1*-*23*, are shown (see Fig. S1A for the whole set of tested temperatures). ***C*)** Synthetic growth defects of a *ten1*-*31 bur1*-*80* double mutant were rescued when either a *ten1*-*31*-*STN1* or a *ten1*-*31*-*CDC13* fusion gene was expressed from a centromeric plasmid under the control of the *TEN1* promoter in the absence of any other form of Ten1 within the cell (rows 1-4). Moreover, a *TEN1*-*STN1* fusion gene rescued *bur1*-*80* at 34°C (compare rows 1 and 5). **(*D*)** *TEN1* geneticaly interacts with *SPT4* and *SPT5*, as well as with *FCP1*, *RPB1* and *CDC73*, as strong synthetic interactions between the corresponding mutations were observed. *ten1*-*31*, *spt5*-*194*, *fcp1*-*1* and *rpo21*-*1* are temperature-sensitive mutations in essential genes, while *spt4*Δ and *cdc73*Δ are null mutations. As in (B), only the most relevant temperatures of growth are shown (see Fig. S3A for the whole set of tested temperatures).

In addition to *ten1*-*33* and *ten1*-*31*, four additional *ten1* mutants also exhibited synthetic interactions with *bur1*-*80* (**Supplemental material** and **Fig. S1B**). Sequencing the *BUR1* genomic locus of *ten1*-*33 bur1*-*27* identified a single point mutation, P281L. The single *bur1*-*27* mutant was found to be temperature-sensitive at 36°C (**Fig. 1A**). This phenotype was not rescued by overexpression of *TEN1* and, vice versa, Ten1 loss of function was not rescued by *BUR1* (**Supplemental material**). Moreover, the observed synthetic lethality between the *ten1* and *bur1* mutations was not due to altered *TEN1* transcription (**Supplemental material** and **Fig. S1D**).

### *CDC13* and *STN1*, unlike *TEN1*, do not genetically interact with *BUR1* and *CAK1*, but yet Cdc13 and Stn1 appear to associate with Ten1 to affect transcription

Based on genetic interactions, Stn1 and Cdc13, unlike Ten1, did not appear to be involved in Bur1-related transcriptional pathways (**Supplementary material** and **Fig. S1C**). However, since Ten1, Stn1 and Cdc13 are known to be together in a complex, we decided to look for further evidence for the implication of the whole CST complex in transcription. To this end we used fusion (hybrid) proteins, a method already applied with success in studies on Cdc13 and Stn1 (Evans and Lundblad 1999; Grandin et al. 2000). First, after expressing in a *bur1*-*80* mutant a Ten1-31-Stn1 fusion protein, we observed that the synthetic growth defects between *bur1*-*80* and *ten1*-*31* were totally suppressed (**Fig. 1C**). A Ten1-31-Cdc13 fusion construct could also rescue the synthetic defect between *bur1*-*80* and *ten1*-*31* (**Fig. 1C**). Most interestingly, expression of a *TEN1*-*STN1* hybrid gene allowed the *bur1*-*80* mutant cells to grow even better than those expressing the *ten1*-*31*-*STN1*/*CDC13* fusions or *TEN1* alone, an effect seen at 34°C (**Fig. 1C**). These experiments indicate that providing a permanent association between either Stn1 or Cdc13 and Ten1-31, by means of expressing hybrid proteins, can eliminate the deleterious effects of the Ten1-31 mutant protein. In addition, providing a permanent association between wild-type Ten1 and Stn1 rescues *bur1*-*80* temperature sensitivity, a situation that is distinct from the synthetic lethality between *ten1*-*31* and *bur1*-*80*. From these experiments, we suggest that Stn1 and Cdc13 most probably cooperate with Ten1 in transcription functions, but that, based on genetics, Ten1 has a more direct and predominant role than those of Stn1 and Cdc13.

### *TEN1* genetically interacts with the RNA pol II transcriptional machinery

In budding yeast, the main role of cyclin-dependent kinase (CDK)-activating kinase (CAK) is the activation, by phosphorylation, of CDKs (**Supplemental material** and **Fig. S2A)**. Cdc28, Kin28, Bur1 and Ctk1, but not Srb10 and Pho85, are phosphorylated by Cak1. Interestingly, the *ten1*-*31* mutant exhibited genetic interactions with *kin28*-*ts*, but not with *srb10*Δ or *pho85*Δ (**Fig. S2B, C** and **data not shown**). On the other hand, *ten1*-*31* exhibited strong synthetic interactions with *fcp1*-*1*, a mutation in the RNA pol II CTD-Ser2P phosphatase, but not with the *rtr1*Δ or *ssu72*-*2* mutations, which inactivates or alters, respectively, RNA pol II CTD-Ser5/7P phosphatases (**Fig. 1D** and **data not shown**; **Fig. S3A**). Using classical genetic methods like that used to construct all double mutants in the present study, namely sporulation of a diploid heterozygous for both genes, we were unable to derive a *ten1*-*31 ctk1*Δ mutant, thus suggesting synthetic lethality between the two mutations. *TEN1* also genetically interacted with the elongation factors-coding *SPT4* and *SPT5* genes, as well as with *CDC73*, coding for a component of the PAF1 transcription elongation complex, and *RPB1*, coding for the largest subunit of RNA pol II (**Fig. 1D** and **Fig. S3A**). All these genetic data strongly suggest a role for Ten1 in RNA pol II transcription in general, but particularly in the elongation step. On the opposite, we did not find any genetic interaction between *ten1* mutants and mutants of the THO complex, indicating that, most probably, Ten1 is not functioning in cooperation with the THO complex to regulate transcription of non coding telomeric DNA into TERRA (see **Supplemental material** and **Fig. S3B, C**).

We next decided to analyze the sensitivity of the *ten1*-*31* mutant to various drugs currently used in the detection of transcription elongation defects such as 6-azauracil (6-AU) (see **Supplemental material** and **Fig. S4A**). Many mutations impairing transcription elongation cause sensitivity to 6-AU, and others, on the opposite, provide resistance to 6-AU, as they constitutively express *IMD2* (Shaw et al. 2001). These particular mutations were found to cause a reduction in the RNA pol II transcription elongation rate (García et al. 2012; Braberg et al. 2013). We found that the *ten1*-*31* mutant was not sensitive to 6-AU (**Fig. S4A**), in agreement with the fact that *IMD2* is constitutively expressed in *ten1*-*31* in the absence of 6-AU and with the fact that *ten1*-*31* suppresses *spt4*Δ sensitivity to 6-AU (**Fig. S4A**). This result suggests again that the *ten1*-*31* mutation may affect transcription elongation (García et al. 2012; Braberg et al. 2013). In addition, the *ten1*-*31* mutant was hypersensitive to formamide, as were *cak1*-*23* and, to a lesser extent, *bur1*-*80* (**Fig. S4B**) (Prelich and Winston, 1993). This drug has been also used to detect mRNA biogenesis defects (Hoyos-Manchado et al. 2017).

### Ten1 influences RNA pol II occupancy during transcription

Next, in order to dissect the role of Ten1 in transcription we performed chromatin immunoprecipitation (ChIP) assays to analyze RNA pol II (Rpb1) association at several regions located within three constitutively transcribed genes (*PMA1*, *YEF3* and *PGK1*) in wild-type and *ten1*-*31* cells at 34°C, a semi-restrictive temperature for mutant growth. We observed a significant decrease in RNA pol II binding to all three genes tested from promoters to the 3’-end regions in *ten1*-*31* when compared to the wild type (**Fig. 2A, E**). Similar to Rpb1, Rpb3 occupancy along the *PMA1* and *PGK1* genes was also reduced in *ten1-31* cells, as in the case of *spt5*-*194* cells (**Fig. S5A**), and this was not due to reduced levels of Rpb1 and Rpb3 proteins (**Fig. S5B**). A slight reduction in RNA pol II binding was also observed in the *stn1*-*154* mutant at the 3’-end of the tested genes (**Supplemental material** and **Fig. S5**).

**Figure 2.**
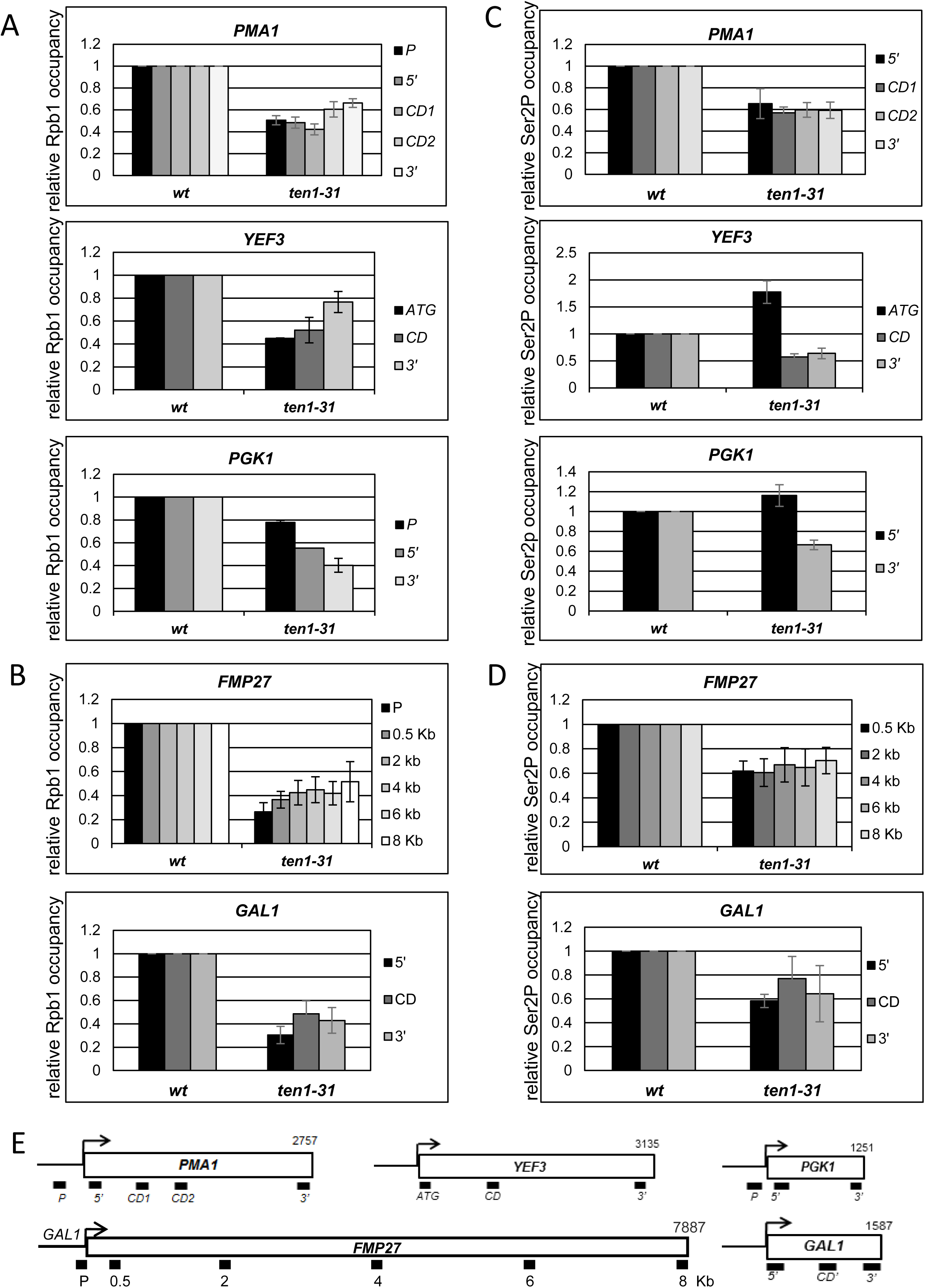
Ten1 influences RNA pol II occupancy during transcription. (***A, B***) Rpb1 gene occupancy is reduced in *ten1*-*31* cells. ChIP analyses were performed in wild-type (*wt*) and *ten1-31* strains grown at 34°C, using an anti-Rpb1 antibody (8W16G). (***A***) Rpb1 occupancy at the promoter (P) or start site (ATG), coding (CD) and 3’-end region (3’) of three constitutively expressed genes, *PMA1*, *YEF3* and *PGK1* were examined by qPCR and quantified (see Materials and methods). Relative Rpb1 binding values obtained in *ten1-31* cells are plotted relative to those from *wt* cells (set equal to 1) for each region. The data plotted here correspond to mean values from at least three independent experiments, and the error bars represent standard errors. (***B***) **Upper panel**: Analysis of Rpb1 occupancy at the promoter (P) and all along coding region of the long gene *FMP27*, expressed under the crontrol of *GAL1* promoter. **Lower panel:** Analysis of *GAL1* gene occupancy by Rpb1 in 5’ coding and 3’-end regions. In both cases the analysis was performed as in (A, B), except that here culturing was done in YPGal medium. (***C***) Levels of Rpb1-Ser2P are altered in *ten1-31* cells. Analysis of Rpb1-CTD Ser2P occupancy at *PMA1*, *YEF3* and *PGK1* genes by ChIP-qPCR using anti-Ser2P antibody (3E-10). Relative Rpb1-Ser2P binding values obtained in *ten1-31* cells are plotted relative to that in *wt* cells (set equal to 1) for each region. The data plotted here correspond to mean values from at least three independent experiments, and the error bars represent standard errors. (***D***) Rpb1-Ser2P occupancy at *FPM27* and *GAL1* genes. The analysis was performed and represented as in (A). (***E***) Schematic representation of the analyzed genes and the position of the primers used for ChIP-qPCR.

We next analyzed Rpb1 distribution along the very long gene *FMP27* (8.0 Kb) whose expression was driven by the rapidly induced *GAL1* promoter as well as along the short *GAL1* gene (1.6 Kb) in the presence of galactose. In *ten1*-*31*, Rpb1 occupancy at the *FMP27* gene was significantly reduced throughout the whole transcription cycle (**Fig. 2B, E**). Rpb1 binding to the transcribed locus was most affected at the promoter, whereas the binding increased in *ten1-31* cells as RNA pol II traveled through the coding region towards the 3’-end. Similar effects were observed for the *GAL1* gene (**Fig. 2B, E**). Therefore, our data suggest that *ten1-31* may affect not only transcription elongation, but also initiation. Altogether, our ChIP data strongly indicate that Ten1 affects RNA pol II association to chromatin during active transcription, thus corroborating our genetic data and pointing out to a role for Ten1 in transcription regulation.

A key mark of the elongation step is the phosphorylation of the Rpb1-CTD Ser2 residues. Thus, Ser2 phosphorylation starts upon promoter clearance and increases all along the transcription cycle until the polymerase reaches the termination region (Buratowski 2003; 2009). Since our data suggest that Ten1 influences transcription elongation, we examined the levels of Rpb1-Ser2P associated to the chromatin during active transcription in *ten1-31* and wild-type cells. As shown in **Figure 2C**, Rbp1-Ser2P binding in *ten1-31* is altered when compared to wild-type cells, in accordance with an elongation defect. Whereas in the *PMA1* gene, Rpb1-Ser2P binding decreases from early elongation (5’) to termination (3’), it is increased at the 5’ region of *YEF3*, and to a lesser extent at that of *PGK1*, though in these two genes, Ser2P binding is reduced in coding and 3’-end regions, similarly to what we observed in the *PMA1* gene. In the case of the galactose inducible genes, *FMP27* and *GAL1*, Rpb1-Ser2P binding is also significantly reduced along the genes (**Fig. 2D**). However, the reduction of Rpb1-Ser2P levels did not appear to be as pronounced as that of Rpb1 levels, suggesting increased Rpb1-Ser2P relative levels. This agrees with Ser2P levels in whole cell extracts being slightly augmented in *ten-31* cells, without changes in Rpb1 levels (**Fig. S5B)**. **Figure S6A, B**, illustrating the ChIP Rpb1-Ser2P/Rpb1 ratios for all tested genes in *ten1*-*31* and the wild type, allows to better appreciate the fact that in the *FMP27* and *GAL1* genes, Rpb1-Ser2P relative levels in *ten1-31* cells were slightly increased all over the coding region and clearly increased in the 5’ region of the *YEF3* and *PGK1* genes. These data suggest that changes in Ser2P profile in *ten1-31* cells may be gene dependent. Moreover, they are consistent with the genetic interaction found between *TEN1* and *BUR1*, because the elongating kinase Bur1 specifically phosphorylates Ser2 near the promoter regions (Qiu et al. 2009). They are also supported by the observation that *TEN1* genetically interacts with *FCP1*, coding for the Rpb1-Ser2P phosphatase. Furthermore, these RNA pol II ChIP data support, once again, a role for Ten1 in regulating transcription elongation.

### Ten1 influences RNA pol II genome-wide distribution

In order to extend our findings and obtain a wider view of *ten1-31* transcription effects, we performed ChIP-seq experiments in which we immunoprecipitated Rpb1 or Rpb1-Ser2P in wild-type (WT) and *ten1-31* cells. As shown in **Figure 3A, B**, the Rpb1 and Ser2P association patterns (IP/INPUT ratios) with protein coding genes in both strains are similar in shape but with different binding values in the case of Rpb1. This was made clearer when the *ten1-31*/wt ratios were represented (**Fig. 3C, D**). The average Rpb1 occupancy profile in the *ten1*-*31* mutant shows a decreased level of binding at 5’ ends, accompanied by an accumulation in the central part of the gene body (**Fig. 3C**) together with an increased presence of Ser2P binding (**Fig. 3D**). In fact, analysis of genes according to their length indicated that the defect in Ser2P phosphorylation is acquired progressively along the coding region, suggesting that Ten1 loss of function provokes an increasing defect on RNA pol II phosphorylation during elongation (**Fig. S7**). Overall, comparison of Rpb1 binding at the *PMA1*, *YEF3* and *PGK1* genes by ChIP-seq (**Fig. S8**) and ChIP-qPCR (**Fig. S6**) indicated similar types of defects.

**Figure 3.**
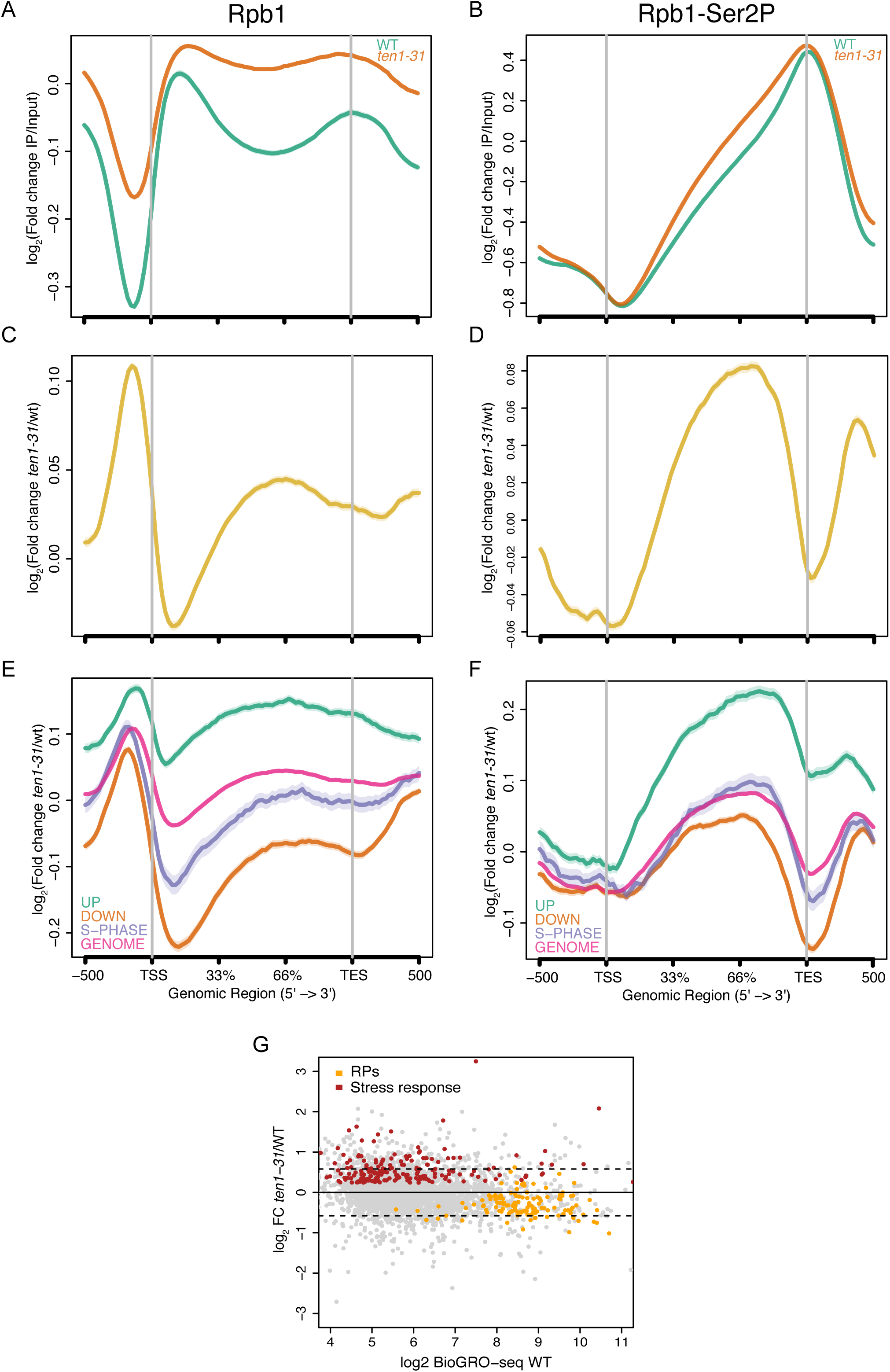
ChIP-seq and RNA-seq analysis of the effects of *TEN1* genetic inactivation. (***A***) Input-normalized average Rpb1 occupancy profile relative to the transcription start site (TSS) and transcription end site (TES) for all annotated protein-coding genes in the yeast genome. The green trace corresponds to the Rpb1 occupancy in the wild-type strain (wt), whereas the orange trace corresponds to the occupancy in the *ten1-31* mutant. The gene body regions have been scaled to an average length and depicted as percentage of the distance from the start, whereas the upstream and downstream flanking regions represent real genomic distances from the TSS and the TES. Normalised occupancy is represented as the log2 Fold Change of Rpb1 IP divided by its corresponding Input. (***B***) Same as in (A) for the Rpb1-Ser2P IP. (***C***) Average differential binding profile for Rpb1-IP in *ten1-31* versus wt. (***D***) Same as (C) for the Rpb1-Ser2P IP. (***E***) Same as in (C) for different gene subgroups compared to the average of the genome. Down are the down-regulated genes found in the differential expression analysis of the transcriptome *ten1-31*/wt (n = 982). Up are the up-regulated genes (n = 980). S phase are differentially expressed genes which have a peak of expression in S phase (n = 282). (***F***) Same as in (E) for the Rpb1-Ser2P IPs. Standard deviations are represented as translucent areas around the solid traces (***G***) MA plot showing the results of the DESeq2 differential expression analysis of the *ten1-31* mutant/wt relative to the mean expression level of each gene in both conditions. Horizontal dashed lines indicate the differential expression cut-off chosen to call genes as up- or down-regulated in our analysis.

Having shown a general defect of the *ten1* mutant in Rpb1 binding all over the genome, we next attempted to evaluate its consequences on global gene expression and performed RNA-seq (**Fig. 3E** and **Fig. S9**). The global transcriptome shows a clear environmental stress response (ESR, Gasch et al. 2000) pattern with Ribosome Protein (RP) genes being downregulated and stress-induced genes upregulated (**Fig. 3G**). Our transcriptomic data indicate that genes that were down-regulated (< 1.5 times) in the *ten1*-*31* mutant exhibited less Rpb1 and Rpb1-Ser2P occupancy, as determined by ChIP-seq (**Fig. 3E, F**), than the average genome level, whereas *ten1*-*31* up-regulated genes (> 1.5 times) had more binding of Rpb1 and Rpb1-Ser2P than average. On the other hand, *ten1*-*31* S phase regulated genes (Santos et al. 2015) were slightly below the average genome level in terms of Rpb1 and Rpb1-Ser2P occupancy.

### Ten1, but also Stn1 and Cdc13, physically and functionally interact with the Spt5 elongation factor

To examine whether Ten1 might have physical partners functioning in transcription mechanisms, we performed mass spectrometry analyses on a strain expressing Ten1-Myc13 at endogenous levels, using anti-Myc antibody. Interestingly, Spt5 was identified in three separate experiments, which was further corroborated by co-immunoprecipitation (co-IP) assays (**Fig. 4A** and **Fig. S10)**. Besides, a physical interaction between Cdc13-Myc13 and Spt5 was also observed (**Fig. 4A** and **Fig. S10**). Moreover, Spt5 was also identified by mass spectrometry as a potential partner of Stn1-Myc13 (**data not shown**). Altogether, these data allow us to conclude that Spt5 might represent a pertinent partner of the CST complex. Confirming these findings, genetic interactions between the temperature-sensitive *spt5*-*194* mutant and the temperature-sensitive *stn1*-*13* and *cdc13*-*1* mutants were observed (**Fig. 4B**). Therefore, the whole CST complex may have a role in transcription elongation, possibly through interactions with Spt5.

**Figure 4.**
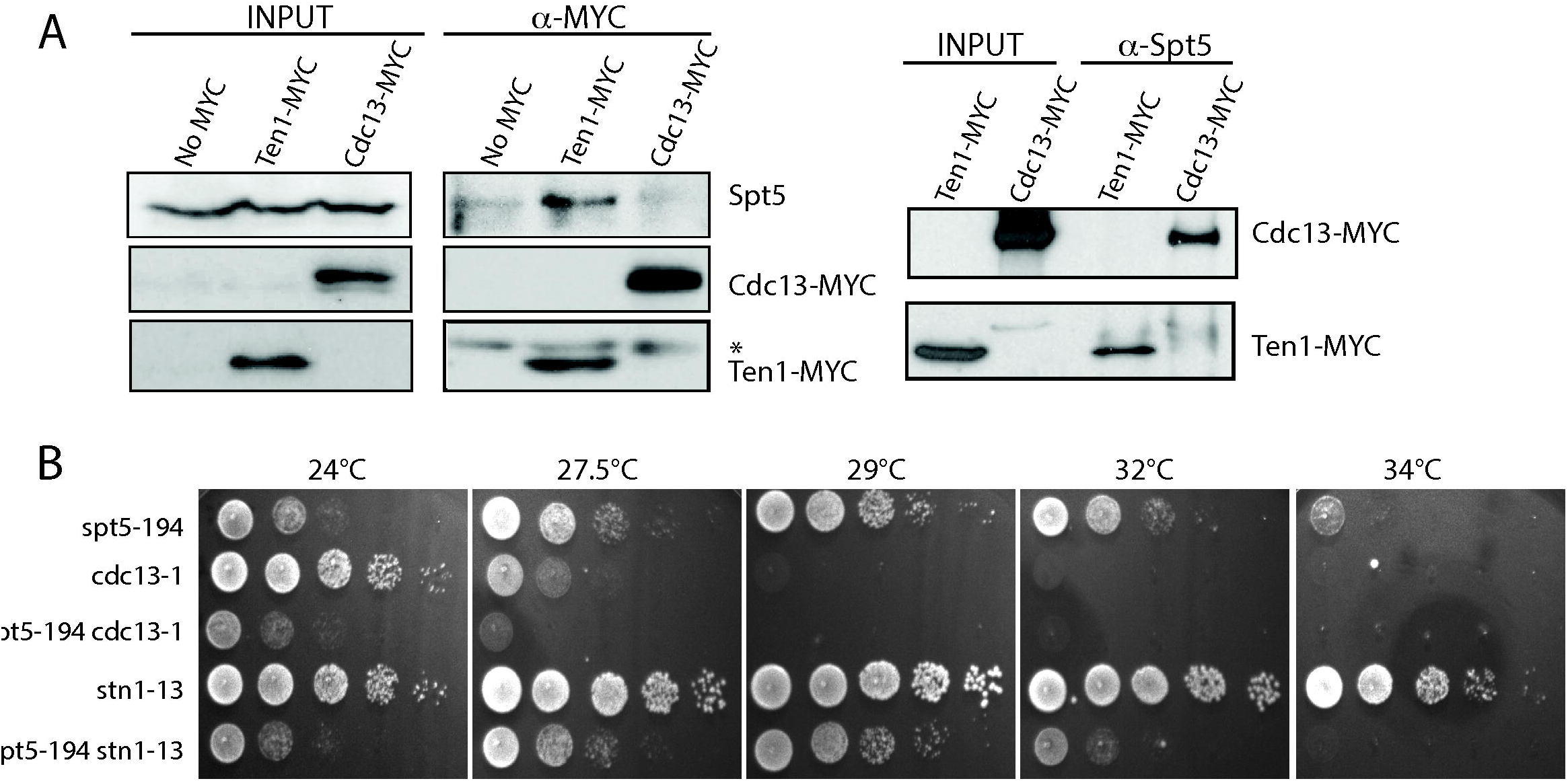
Ten1, but also Stn1 and Cdc13, physically and functionally interact with the Spt5 elongation factor. (***A***) Ten1-Myc_13_ physically associates with Spt5 by co-IP. Cell extracts from asynchronous wild-type cells harboring either endogenous copy of Ten1-Myc_13_ (Holstein et al. 2014) or of Cdc13-Myc_13_ (Oza et al. 2009) were immunoprecipitated with either anti-Myc (**left panel**) or anti-Spt5 (**right panel**) antibody. Input and IPs were analyzed by western blotting with antibodies against the indicated proteins. (* non specific band) (***B***) Both *CDC13* and *STN1* exhibit genetic interactions with *SPT5*. The *spt5*-*194*, *stn1*-*13* and *cdc13*-*1* temperature-sensitive mutations were combined together and growth of double mutants compared with that of each single mutant at the indicated temperatures.

### Ten1 is important to maintain proper levels of Spt5 associated to chromatin during active transcription

Next, we performed ChIP assays in *ten1*-*31* and wild-type cells expressing Spt5-Flag to further investigate the interactions between CST and Spt5 (**Fig. 5**). Spt5-Flag occupancy was significantly reduced in *ten1-31* cells compared with the wild type for all five tested genes (**Fig. 5A, C**), while Spt5 protein levels remained unchanged (**Fig. 5B**). It is worth mentioning that in the case of the extra long *FMP27* gene, and to lesser extent in the long *YEF3* gene, we clearly observed an increase of Spt5-Flag occupancy from the 5’-end to the 3’-end regions. This association pattern was similar to that observed above for Rpb1, and comparable to those previously observed in elongation rate mutants (Quan and Hartzog 2010; García et al. 2012). This also correlates with *ten1-31* cells being resistant to 6-AU treatment (Fig. S4), likewise to some *rpb1* and transcription factors mutants in which RNA pol II transcription elongation rate is slowed down (García et al. 2012; Braberg et al. 2013). Therefore, our results clearly suggest that in *ten1*-*31* cells the elongation rate is reduced. Moreover, our findings are supported by the genetic interactions between *ten1* and *bur1* mutants because Bur1, not only phosphorylates Rpb1-CTD (Qiu et al. 2009), but also regulates the activity of Spt5 by phosphorylation, thus promoting transcription elongation (Liu et al. 2009).

**Figure 5.**
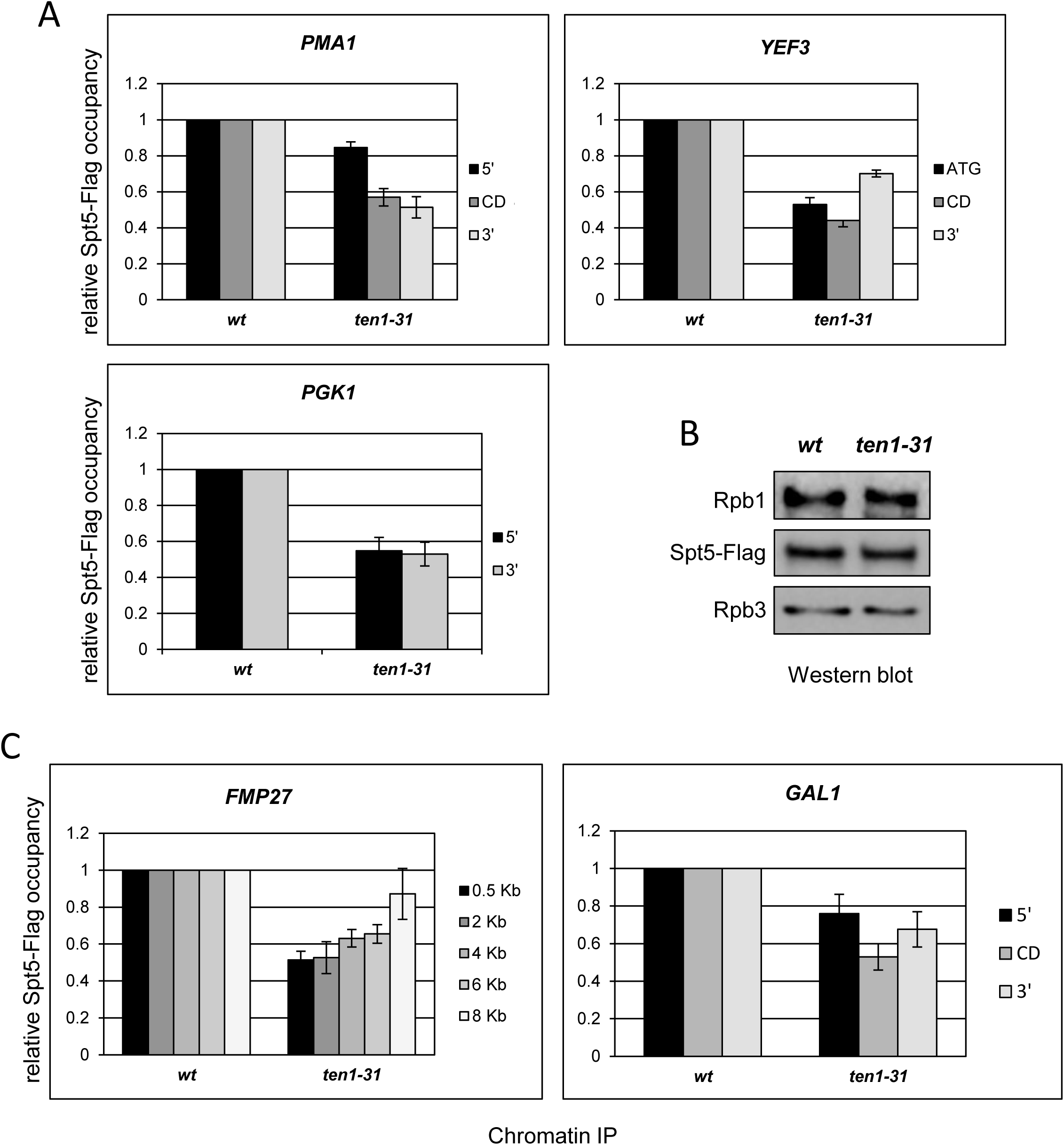
Ten1 is important to maintain proper levels of Spt5 associated to chromatin during active transcription. ChIP analyses were performed using wild-type (*wt*) and *ten1-31* strains grown at 34°C, either in a medium containing glucose to analyze Spt5-Flag binding to the constitutively expressed genes, *PMA1*, *YEF3* and *PGK1* (***A***) or in a medium containing galactose to analyze Spt5-Flag binding to the inducible *GAL1*-*FMP27* and *GAL1* genes (***C***). Spt5-Flag binding was examined by qPCR and represented as in Figure 2. (***B***) Spt5 total protein levels are not altered in *ten1*-*31* mutant cells. Levels of Spt5-Flag were analyzed by western blotting using WCE from wild-type (*wt*) and *ten1-31* cells expressing Spt5-Flag. Levels of Rpb1 and Rpb3 were also tested. Shadows along the curves mean to represent standard deviations.

### The high-mobility group box (HMGB) protein Hmo1 binds Ten1 and genes down-regulated in *ten1-31* are preferentially bound by Hmo1

Besides Spt5, our mass spectrometry experiments identified two proteins that were isolated four times: in all three different experiments using Ten1-Myc13 as the bait and in one experiment using Stn1-Myc13 as the bait. These proteins are Hmo1 and Nhp6B and both have been previously implicated in transcription regulation (Travers 2003; Panday and Grove 2017). The interaction between Ten1-Myc13 and Hmo1-HA2 was confirmed by co-IP (**Fig. 6A**). Since, as mentioned above for *ctk1*Δ, numerous attempts to derive a *ten1*-*31 hmo1*Δ double mutant failed, it is possible that the functional interactions between Ten1 and Hmo1 are so strong that loss of function of both cannot be tolerated by the cell.

**Figure 6.**
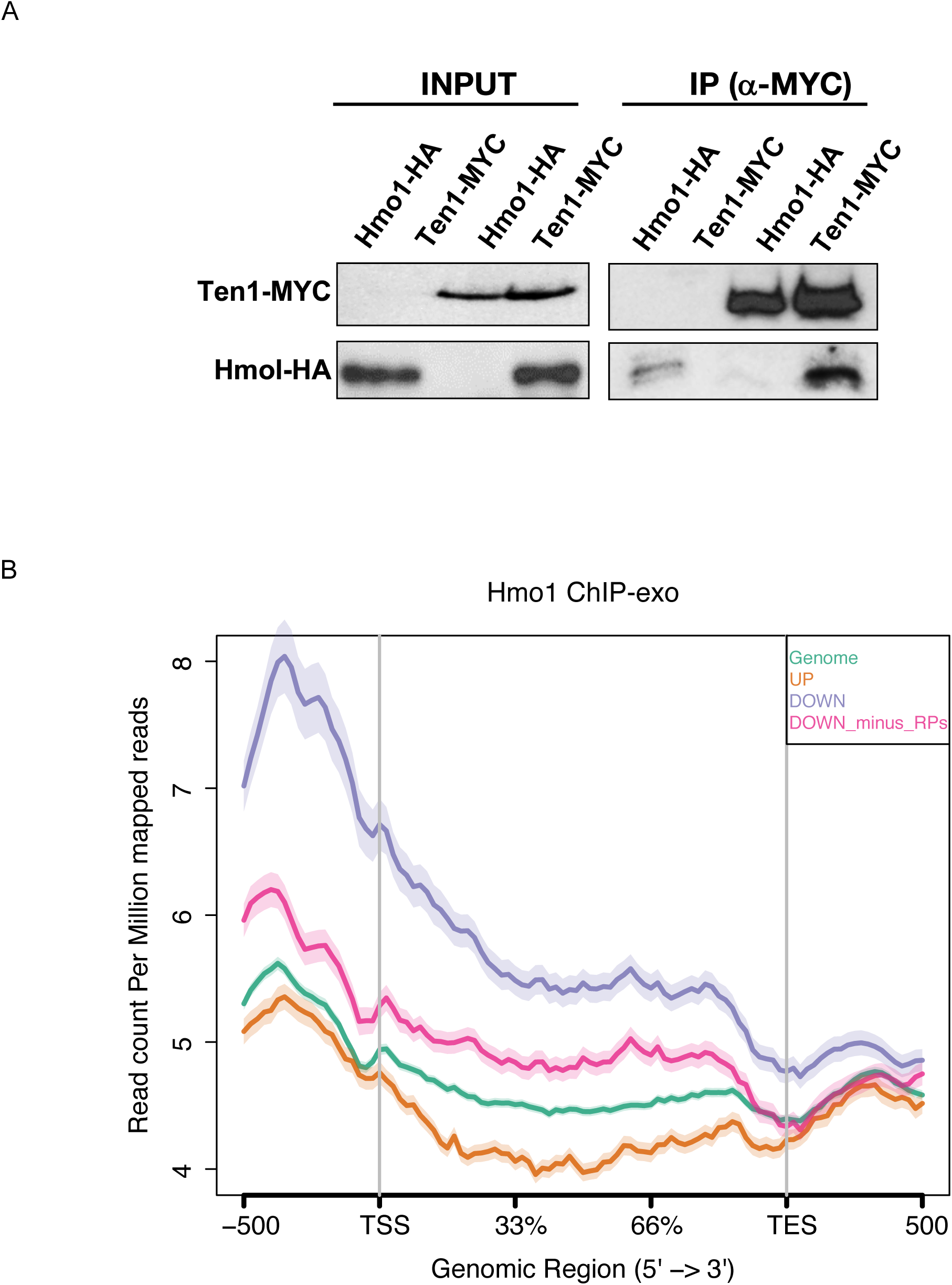
Hmo1 interacts with Ten1 and binds to promoters and gene bodies of genes silenced upon *TEN1* genetic inactivation. **(A)** Ten1-Myc_13_ associates with Hmo1-HA_2_ as determined by co-IP. The assay was performed with WCEs from Hmo1-HA_2_, Ten1-Myc_13_, and Hmo1-HA_2_ Ten1-Myc_13_ cells using an anti-Myc antibody. Inputs and IPs were analyzed by western blotting with antibodies directed against the indicated proteins, as indicated. (***B***) ChIP-exo Hmo1 occupancy profile relative to the TSS and pA sites for different subsets of genes compared to the average of the genome. Down are the down-regulated genes found in the differential expression analysis of the transcriptome *ten1-31*/wt (n = 982). Down_minus_RPs are the down-regulated genes after removing ribosomal protein genes (n = 899). Up are the up-regulated genes (n = 980). S Phase are genes with a peak of expression in the S phase of the cell cycle (n = 877). Standard deviations are represented as translucent areas around the solid traces.

At the whole genome level, Hmo1 tends to accumulate at promoter regions of genes to which it associates (Reja et al. 2015). Therefore, we wondered whether this pattern could be altered in *ten1-31* cells, as potentially suggested by the existence of a Ten1-Hmo1 physical association. Interestingly, most of the *ten1*-*31* down-regulated genes showed the highest Hmo1 occupancy, while up-regulated genes were on average less bound by Hmo1 than at genome-wide level (**Fig. 6B**). Even when we subtracted RP genes from the list of *ten1-31* down-regulated genes **(Fig. 3G)**, we still observed an important defect in Hmo1 binding, therefore indicating a general effect. These data suggest that in the *ten1*-*31* mutant, the expression of Hmo1-bound genes genes is affected because they lack the activating effect of Ten1.

### Genes that are differentially expressed in S phase in *ten1-31* bind less RNA pol II than the rest of the genome

Hmo1 was found to be preferentially recruited at Top2-bound regions of S phase-arrested cells, principally, but not exclusively, at gene promoters. Hmo1 and Top2 were proposed to prevent damage at sites of S phase transcription upon collision with an incoming replication fork (Bermejo et al, 2009). Analyis of Hmo1 ChIP-exo data from Reja et al. (2015) revealed that S phase genes have a higher average Hmo1 occupancy than the rest of the genome. However, such occupancy level is lower than that of down-regulated genes in our *ten1-31* transcriptome analysis **(Fig. 6B)**. Given the genome-wide correlation, established above, between Hmo1 occupancy and *ten1*-*31* differentially expressed genes, we next re-examined our RNA-seq and ChIP-seq data in more detail. First, we found that among the 877 genes that have a peak of high expression in S phase (Santos et al. 2015), there is a statistically significant enrichment (p-value 9 10^-4^) of 282 genes down-regulated in the *ten1*-*31* mutant. Most interestingly, in *ten1*-*31*, down-regulated S phase genes were specifically less bound by RNA pol II overall than the rest of the genome, though accompanied by an increase in Ser2P binding (**Fig. 3F**). Therefore, we conclude that the population of *ten1*-*31* down-regulated genes that preferentially bind Hmo1 at their promoters exhibits a deficit in RNA pol II occupancy, compared to the rest of the genome. Again, all these data together support the hypothesis that Ten1 influences transcription elongation.

### *MRC1* and *CTF18*, coding for DNA replication damage sensing proteins, genetically interact with *TEN1* and *STN1*

Because of the findings above (summarized in **Fig. 7A**), we wondered whether there might be a specific context where CST might bind and affect RNA pol II. Human STN1 and CTC1 were first isolated in biochemical experiments as alpha accessory factors, AAF-44 and AAF-132, respectively, of the DNA polymerase α complex (Goulian et al. 1990; Casteel et al. 2009). Although it has been known for some time that Cdc13 physically interacts with Pol1, the largest subunit of the DNA pol α complex (Qi and Zakian 2000), and Stn1 with Pol12, a subunit of DNA pol α (Grossi et al. 2004), it is actually unknown where these interactions take place exactly (at the telomeres or at the replication forks or both?). Ctf18 and Mrc1 are, together with Mec1 and Rad53, the most important actors in maintaining replication fork integrity upon DNA replication stress (Crabbé et al. 2010). We speculated that CST is normally present at the replication fork and that potential collisions between the moving fork and an incoming transcription unit might activate these DNA replication checkpoint proteins. If CST responds to such a stress to affect transcription accordingly, then mutations in CST might be synthetically lethal with mutations in *MRC1* or *CTF18*. Indeed, strong genetic interactions between *CTF18* and *TEN1*, *CTF18* and *STN1*, as well as between *MRC1* and *TEN1*, were observed (**Fig. 7B**). These results suggest that CST functions in transcription regulation might take place at the replication fork.

**Figure 7.**
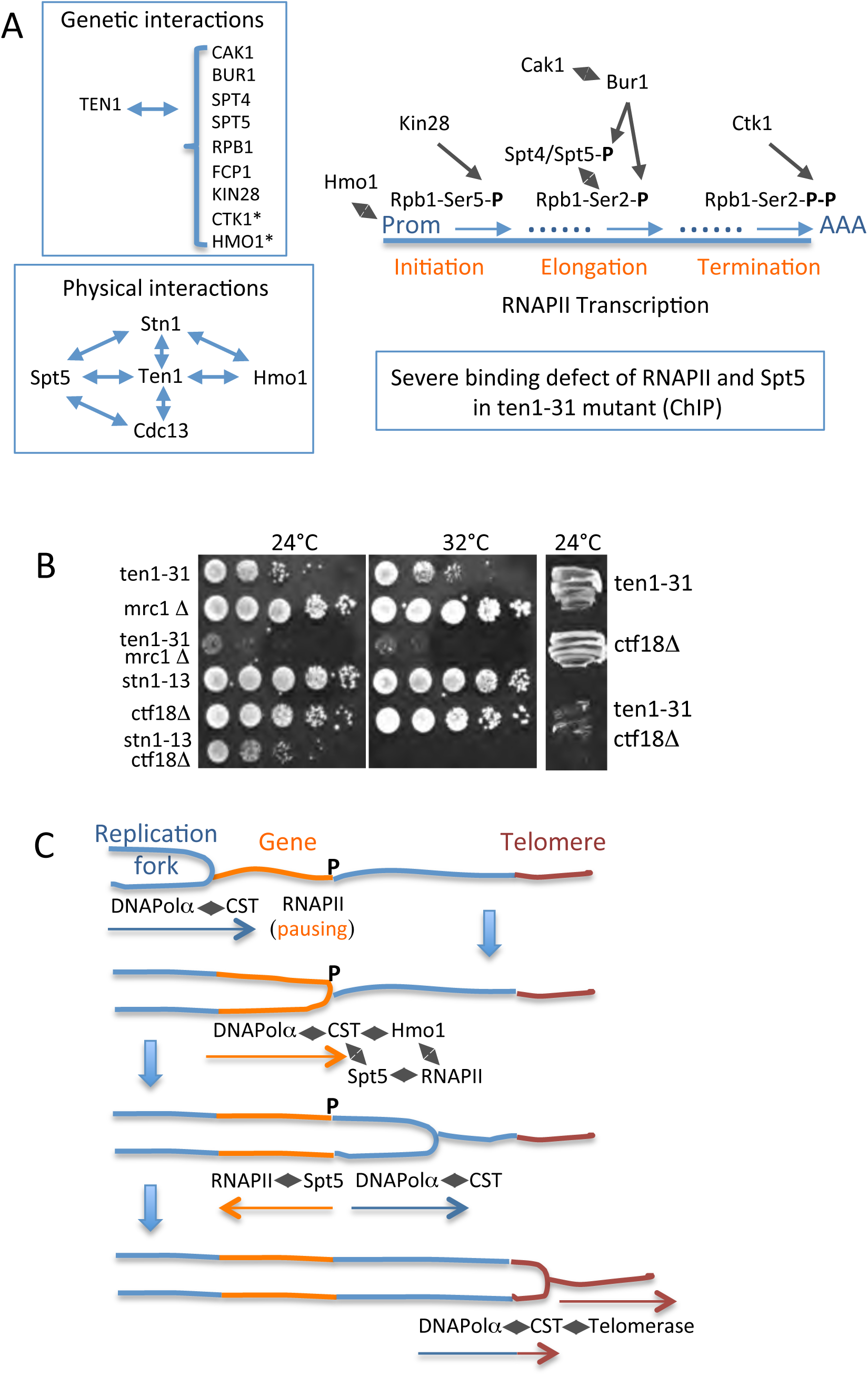
(***A***) Schematic overview of the data presented in this work supporting a role for CST in RNA pol II transcription. Asterisks indicate situations in which the double mutants could not be derived, most probably because of complete inviability. Black diamonds represents physical interactions (***B***) Genetic interactions between mutations in CST components, *ten1*-*31* and *stn1*-*13*, and null mutations in the DNA replication stress checkpoints Mrc1 and Ctf18. (***C***) Hypothetical working model for a role for the Cdc13-Stn1-Ten1 complex at transcribed genes encountering progressing replication forks. We speculate that, at the replication fork, Cdc13 and Stn1, which physically associate with Pol1 and Pol12 (Pol α), respectively, perceive a signal emitted by the Mec1, Rad53, Ctf18 and Mrc1 DNA replication checkpoints, which themselves have sensed mechanical vibrations resulting from the clash between the replication fork and the transcription machinery moving in opposite directions. This signal provokes the dissociation between CST and Pol α, allowing the CST complex to attach the promoter of the transcribing gene, via Hmo1, before activating the RNA pol II machinery, via Spt5 (see main text for detail).

## Discussion

### Ten1 has a function in transcription elongation in association with Stn1 and Cdc13

Several sets of data demonstrate that Ten1 functions in regulating RNA pol II transcription in association with Stn1 and Cdc13, these three proteins forming the essential *S*. *cerevisiae* telomeric CST complex. First, Ten1 influences association of Spt5, a major, highly conserved, player in transcription elongation, with actively transcribed chromatin, as well as RNA pol II distribution during the whole transcription cycle. We also showed that Cdc13 and Stn1 physically associate with Spt5 and genetically interact with it. Second, Rpb1 and Rpb1-Ser2P levels, which are crucial to correctly maintain transcription elongation, are also altered in the *ten1* mutant. Consistent with this, *ten1-31* genetically interacts with the CTD-Ser2P phosphatase, Fcp1, and the CTD-Ser2P kinase, Bur1, as well as with the Spt4/5 and Cdc73 elongation factors. Besides, *ten1-31* displayed 6-AU resistance, a feature common to mutations causing a decrease in elongation rate (Mason and Struhl 2005; García et al. 2012; Braberg et al. 2013). Third, our experiments with the *ten1*-*31*-*STN1* and *ten1*-*31*-*CDC13* hybrid genes suggest that all three components are acting together to regulate transcription. Actually, the *stn1*-*154* mutant conferred mild defects in the association of RNA pol II with the 3’-end region of long genes. Finally, the CST complex seems to specifically participate in RNA pol II transcription, as we have not observed that the *ten1*-*31* mutation affects at least RNA pol I association to the rDNA gene (**Fig. S11**).

### CST has a potential role in stimulating the transcription machinery upon collision with a replication fork

Our working model starts with the likely possibility that CST action on transcription may be initiated at the progressing replication fork (**Fig. 7C**). We speculate that, upon torsional stress provoked by the imminent arrival of a moving transcription unit in front of the progressing replication fork, the checkpoint sensors Ctf18 and Mrc1 (and also Mec1 and Rad53, the effectors of both sensors) arrest the progression of the replication fork (among other events). A transient dissociation between Cdc13 and Pol1 and/or Stn1 and Pol12, both components of the DNA pol α complex then allows CST to move towards the colliding transcription unit and establish contacts with Hmo1 (**Fig. 7C**). Supporting this hypothesis, we have found physical interactions between Pol1 and Spt5 by two-hybrid (**Fig. S12**), thereby confirming a similar interaction detected by mass spectrometry (Lindstrom et al. 2003).

Recent findings have established that in S phase-arrested cells, Hmo1 was preferentially recruited at Top2-bound regions, principally at gene promoters. This led to the proposal that Top2 (and also Top1) and Hmo1 might solve difficult topological contexts in S phase when transcription has to face incoming replication forks (Bermejo et al. 2009). Top1, Top2 and Hmo1 (Bermejo et al. 2009), together with Sen1 (Alzu et al. 2012), appear to be sufficient to manage head-on collisions between the transcription and replication machineries. We propose, based on our finding that Hmo1 occupancy is higher than average at *ten1*-*31* down-regulated genes, that the CST complex might play an important role in stimulating the RNA pol II machinery, principally through physical interactions with Spt5, after it has collided with the replication fork or, at least, in synchronizing both machineries (**Fig. 7C**). Interestingly, the *ten1*-*31 hmo1*Δ double mutant was inviable and the *ten1*-*31* mutant exhibited synthetic interactions with the *top1*Δ *top2*-*1* double mutant (**Fig. S13**), thereby suggesting the existence of functional interactions between Ten1 and the three proteins implicated in managing transcription/replication collisions (Bermejo et al. 2009).

Budding yeast CST might also stimulate DNA pol α activity after the collision, as proposed for mammalian CST in face of a DNA replication stress (Stewart et al. 2012; Kasbek et al. 2013). In our model, yeast CST travels with the replication forks and arrives at the extremities of the telomeres at the right time, during late S phase, to occupy the elongating single-stranded G-overhang (Wellinger et al. 1993). This way, CST having accomplished its functions of transcriptional regulation during S phase executes its telomeric functions immediately after (**Fig. 7C**).

The situation with Ten1 described here bears striking resemblances with that concerning Hog1. Indeed, upon osmostress, the Hog1 MAP kinase interacts with components of the RNA pol II transcription elongation complex such as Spt4, Paf1, Dst1 and Thp1 to recruit the RSC chromatin remodeler complex to stress-responsive genes (Mas et al. 2009; Nadal-Ribelles et al. 2012). Bearing similarities with this situation, Ten1 is functionally linked to Spt5 and genetically linked to Cdc73, a subunit of the PAF1 complex (Liu et al. 2009). Therefore, the Spt4/5 and PAF1 complexes might represent a privileged location within the RNA pol II transcription machinery to regulate transcription upon either external stress or stress provoked by fork progression.

Other telomeric proteins are also known to play a role in transcription. For instance, in mouse embryonic cells and human cancer cells, RAP1 and TRF2 endorse extratelomeric functions and are true regulators of transcription (Martinez et al. 2010; Yang et al. 2011; Ye et al. 2014). *S*. *cerevisiae* Rap1, another telomeric protein, is also a true transcription factor as it modulates expression of many genes, including ribosomal protein genes, MATα genes, several glycolytic enzyme genes (Buchman et al. 1988), as well as genes that adapt chromatin changes in response to telomeric senescence (Platt et al. 2013).

Intriguinly, several *S*. *cerevisiae* mutants of factors involved in RNA pol II transcript biogenesis have been found to exhibit altered telomere length (Ungar et al. 2009). It has been argued that, since telomerase and most of its regulators are present in the cell at extremely low amounts, even slight changes in transcription regulators might significantly affect telomere length (Ungar et al. 2009). Alternatively, these transcription mutants might necessitate increased levels of CST proteins to manage head-on collisions, ending up in a deficit of telomeric CST, thereby affecting telomere length.

In summary, the present data uncover a completely novel facet of the telomeric Cdc13-Stn1-Ten1 complex, namely a role in the regulation of transcription, potentially serving to optimize the functioning of the RNA pol II machinery upon head-on collision with a replication fork, following signaling by Hmo1. Noticeably, in our model, CST might also be in charge for coordinating the completion of S phase with the onset of telomere replication by telomerase/Pol α. Therefore, the CST complex now appears as a versatile machine with several distinct functions that take advantage of the properties of each of its three components at different times of the cell cycle and are based on several different protein-protein interactions, the principal ones being those with Pol1, Pol12, Est1 and, as shown here, Spt5, as well as on the ssDNA-binding properties of Cdc13 and Stn1. Additionally, a well established role of Spt5 is to release paused or arrested RNA pol II and promote transcription elongation in higher eucaryotes (Hartzog and Fu 2013). Therefore, based on our data, it is possible that Spt5 might also be acting to promote the release of paused or arrested RNA pol II from sites where the transcription and replication machineries are prone to collide. The present finding of the existence of extra-telomeric functions for Ten1 in the regulation of RNA polymerase II in cooperation with Stn1 and Cdc13 has profound repercussions on future studies both on telomeric and transcription pathways.

## Supporting information

## Star methods

**Detailed methods are provided in the online version of this paper and include the following:**

Yeast strains and media

Genetic screen to find extragenic mutations enhancing the *ten1* phenotype

Co-immunoprecipitation and western blot analysis

Two-hybrid experiments

RNA isolation and RT-PCR

RNA-seq

## Accession numbers

Raw and processed data are available at GEO under the accession number GSE120296.

Chromatin immunoprecipitation (ChIP) and ChIP-seq

Mass spectrometry analysis

## Supplemental material

Supplemental material is available online.

## Acknowledgments

We are grateful to Alberto Paradela and Adán Alpízar Morúa from the Proteomic Department at the CNB, CSIC Madrid, Spain, for help with the mass spectrometry experiments. We thank people at the Genomic Facility at the CRG in Barcelona. We also thank David Lydall, Craig Peterson and Rolf Sternglanz for the gift of strains. This work was supported by grants from the “Fondation de France” and from the “Ligue Grand-Ouest contre le Cancer” to MC, from the Consejo Superior de Investigaciones Cientificas (i-LINK1213) to OC and MC, and from the Spanish Ministry of Economy and Competitiveness (MINECO) (BFU2017-84694-P) to OC and to JE.P-O. (BFU2016-77728-C3-3-P) and from Generalitat Valenciana to JE.P-O. (PROMETEOII 2015/006). We also thank the Spanish Excellence Network *RNA life* (BFU2015-71978-REDT).

## Authors contributions

NG, OC and MC conceived and designed the experiments. NG, NGP, EM, OC and MC performed the experiments. AJ and JEPO analyzed RNA-seq and ChIP-seq data. NG, OC and MC analyzed the data. NG, OC and MC wrote the manuscript. All authors revised and approved the manuscript.

