## Supplementary material for "The telomeric Cdc13-Stn1-Ten1 complex regulates RNA polymerase II transcription"

**SUPPLEMENTAL MATERIAL**

**Supplemental results**

*Isolation of a mutant of* SGV1*/*BUR1 *as exhibiting synthetic growth defects with ten1 mutants*

After comparing growth of ~ 40,000 colonies of mutagenized *ten1*-*16*, *ten1*-*31* and *ten1*-*33* mutant strains at 24°C *versus* 36°C, as described in Materials and methods, 17 mutants were initially selected on the basis of exhibiting aggravated growth at 36°C. These 17 mutants were next transformed with a 2μ plasmid overexpressing *TEN1*. Double mutant strains whose growth was not improved by *TEN1* overexpression, namely 7/17, were discarded, because this pointed out to a possible lack of specificity of the observed genetic interaction. The 10 remaining strains were then back-crossed with the original *ten1* mutant to see whether 2:2 segregation of the phenotype at 36°C took place, thereby indicating that the observed phenotype likely resulted from a single mutation. Cloning of the *ten1* extragenic mutations was then attempted through complementation experiments with YCp50 or YEp24 genomic DNA libraries (centromeric and episomal, respectively), both bearing a selectable *URA3* marker (Carlson and Botstein 1982; Rose et al. 1987). From those 10 mutants, only one could be successfully complemented by a genomic library DNA fragment that suppressed the aggravated growth arrest at 36°C. Thus, the so-called *ten1*-*33* mut. #27 double mutant could be complemented by a YEp24 library (2μ, episomal) fragment of ~ 8.3 Kb corresponding to a region of chromosome XVI comprised between nt #867,095 and nt #858,757. Further genetic analyses identified *SGV1*/*BUR1* (YPR161C), from now on referred to as *BUR1*, as the gene rescuing the genetic interaction between mut. #27 and *ten1*-*33*. In the same complementation experiment, *CAK1* (YFL029C) was also isolated as complementing the loss of function of *ten1*-*33* mut. #27 (after further restricting the YEp24 genomic fragment of ~ 10.1 Kb corresponding to nt #72,543 to nt #82,623 of chromosome VI).

Since *BUR1* and *CAK1* overexpression could both rescue the phenotype of the *ten1*-*33* mut. #27 double mutant, additional genetic analyses were performed to find out whether one of these genes was responsible for the phenotype of the double mutant when mutated and, if yes, which one. To this end, the original *ten1*-*33* mut. #27 double mutant was backcrossed against either the temperature-sensitive *bur1*-*80* mutant (Keogh et al. 2003) or the temperature-sensitive *cak1*-*23* mutant (Espinoza et al. 1998). These experiments showed that mutation #27 could never segregate with *bur1*-*80*, while, on the other hand, this mutation could segregate with *cak1*-*23*, indicating that the *BUR1* locus was mutated in the *ten1*-*33* mut. #27 double mutant. Therefore, these genetic analyses established that *CAK1* was only acting as a suppressor of *ten1*-*33* mut. #27 (from now on referred to as *bur1*-*27*), in agreement with the previous finding that 2μ overexpression of *CAK1* could rescue the temperature sensitivity of the *bur1*-*1* mutant (Yao and Prelich 2002). As expected from these findings, overexpression of *BUR1*, but not of *CAK1*, from a centromeric (CEN) plasmid also rescued the aggravated defect of *ten1*-*33* *bur1*-*27* at 36°C (**data not shown**).

The temperature-sensitive *bur1*-*27*, *bur1*-*80* and *cak1*-*23* mutants were not rescued by CEN or 2μ overexpression of *TEN1* (**data not shown**). In addition, neither 2μ overexpression of *BUR1* nor *CAK1* rescued the growth defects of the temperature-sensitive *ten1*-*31* and *ten1*-*33* mutants (**data not shown**). As expected, *bur1*-*27* expressed from a CEN plasmid failed to rescue the *ten1*-*33 bur1*-*27* double mutant at 36°C (**data not shown**).

The temperature-sensitive *bur1*-*80* and *cak1*-*23* mutants also exhibited synthetic growth defects with *ten1*-*31* (**Fig. 1B** and **Fig. S1A**). Note that the *ten1*-*31* *cak1*-*23* double mutant could be rescued, at 29°C, by overexpression (2μ multicopy) of *CAK1* but not by overexpression of *BUR1* (**data not shown**). Knowing that YEp-*CAK1* also rescued *ten1*-*33* *bur1*-*27* at 36°C, this tends to suggest that *cak1*-*23* might cause additional defects in combination with *ten1*-*31* independently from *BUR1*. In addition to *ten1*-*31* and *ten1*-*33*, two other temperature-sensitive *ten1* mutants, *ten1*-*16* and *ten1*-*100*, as well as two non temperature-sensitive *ten1* mutants, *ten1*-*3* and *ten1*-*6*, previously isolated on the basis of harboring elongated telomeres (Grandin et al. 2001) also exhibited synthetic growth defects with *bur1*-*80* (**Fig. S1B**).

*Absence of genetic interaction between the other* CST *components and* BUR1-CAK1

Previous data have established that Cdc13, Stn1 and Ten1 physically interact together either by two hybrid or co-immunoprecipitation (Grandin et al. 1997; 2001). Although these proteins are frequently referred to as the CST complex, there has been evidence suggesting that Cdc13 and Stn1 performed overlapping but distinct functions. At this point, we wanted to know whether there were also genetic interactions between *BUR1* or *CAK1* and *STN1* or *CDC13*. No genetic interactions were observed between *cdc13*-*1* and *bur1*-*80* or *cak1*-*23,* nor between *stn1*-*13* and *bur1*-*80* or *cak1*-*23* (**Fig. S1C**). In addition, *stn1*-*101*, *stn1*-*138* and *stn1*-*154*, all three temperature-sensitive mutants of *STN1* (Grandin and Charbonneau 2001; Grandin et al. 2001) did not exhibit any synthetic growth defects with *bur1*-*80* (**Fig. S1C**).

Since Bur1 and Cak1 function in transcription, one could argue that the reason for the synthetic growth defects between the *ten1* and *bur1* or *cak1* mutations might be due specifically to alterations in *TEN1* transcription (but not in *STN1* and *CDC13* transcription) provoked by the knock-out of Bur1-controlled transcriptional pathways. To eliminate this possibility, the *ten1*-*33* allele was placed under the control of *STN1*’s natural promoter. Since *stn1* mutations are not synthetic with a mutation in *BUR1*, as seen above, we inferred that mutations in Bur1-related pathways did not result in a significant defect in *STN1* transcription. Yet, *ten1*-*33* ORF under the control of *STN1*’s promoter still exhibited a synthetic interaction with *bur1*-*80* (**Fig. S1D**).

TEN1 *genetically interacts with various genes coding for transcriptional regulators, including* RPB1 *coding for the largest subunit of RNA pol II*

CDKs, initially discovered as major regulators of cell cycle transitions, were later recognized as conserved regulators of RNA pol II transcription. In *S*. *cerevisiae* and mammals, Bur1/CDK9 and Ctk1/CDK12 act in transcription elongation, Kin28/CDK7 during transcription initiation and Srb10/Ssn3/CDK8 is a Mediator subunit proposed to inactivate RNA pol II prior to pre-initiation complex formation (Hengartner et al. 1998). Among the CDKs, Pho85 does not seem to play a role in transcription, but rather in phosphate metabolism and cell cycle control (Jiménez et al. 2013), while Cdc28, which indirectly activates particular transcription programs, appears to also directly stimulate transcription of a subset of housekeeping genes (Chymkowitch and Enserink 2013).

After uncovering the *TEN1*-*CAK1* genetic interactions, we asked whether, like *BUR1*, genes coding for the other CDKs also exhibited a genetic interaction with *TEN1*. *ten1*-*31* exhibited synthetic growth defects with a temperature-sensitive *kin28* mutation (**Fig. S2C**), but not with *srb10Δ* (**Fig. S2B**) or *pho85*Δ . Interestingly, *bur1* and *bur2* mutants were previously found to exhibit synthetic growth defects with *ctk1*Δ, but not with *srb10*Δ or temperature-sensitive *kin28* mutants (Lindstrom and Hartzog 2001; Murray et al. 2001). **Fig. S3A** shows in more detail the genetic interactions of *ten1*-*31* with transcription regulators mutants illustrated in **Figure 1D**.

*Ten1 does not appear to function in telomeric DNA transcription (TERRA)*

The THO complex, which promotes efficient packaging of nascent mRNAs into ribonucleoprotein complexes, has recently been shown, in *S*. *cerevisiae*, to be a component of telomeres and implicated in TERRA regulation (Pfeiffer et al. 2013; Azzalin and Lingner 2014). Because Ten1 is a telomeric protein, we wondered whether its implication in transcription, described above, might be correlated with the implication of THO in the transcription of telomeric DNA into TERRA. Mutations in components of the THO complex, namely *mft1Δ* and *thp2Δ*, did not aggravate *ten1*-*31* growth defects (**Fig. S3B**). Mutants of the THO complex accumulate RNA:DNA hybrids or R-loops, which are routinely removed by the RNase H nucleases, Rnh1 and Rnh201. Null mutations in either one of these two Rnase H nucleases or in both did not exhibit any particular phenotype or growth defects when combined with the *ten1*-*31* mutation (**Fig. S3C**). Finally, we found that *ten1*-*31* growth defects were not suppressed by overexpression of *RNH1*, under the control of the inducible *GAL1*/*10* promoter (**data not shown**). Therefore, it is unlikely that accumulation of R-loops (which also takes place in some mutants of transcription termination, *sen1* mutants for instance) is responsible for the phenotype of *ten1*-*31*.

*Sensitivity of* ten1-31 *to drugs affecting transcription elongation*

Some *BUR1* mutants display hypersensitivity to the nucleotide analog 6-azauracil (6-AU) and mycophenolic acid (MPA) (Murray et al. 2001; Keogh et al. 2003). Both drugs have been shown to affect the rate of transcription elongation *in vivo* by depressing cellular GTP levels, which induces the transcription of the *IMD2* gene in wild-type cells (Exinger and Lacroute 1992). Many mutations impairing transcription elongation cause sensitivity to 6-AU or MPA treatment, while others, on the opposite, provide resistance to these drugs, as they constitutively express *IMD2* (Shaw et al. 2001). We found that the *ten1*-*31* mutant was not sensitive to 6-AU, contrary to the *spt4*Δ mutant, but similar to *spt5*-*194* (**Fig. S4A**), and was not sensitive either to 15 μg/ml MPA (**not shown**). This was in agreement with the facts that *IMD2* is constitutively expressed in *ten1*-*31* in the absence of 6-AU (**Fig. S4A**) and that *ten1*-*31* suppressed *spt4*Δ sensitivity to 6-AU, although it clearly increased the growth defect of *spt4*Δ at 29°C (**Fig. S4A**). Most interestingly, 6-AU slightly suppressed the temperature sensitivity of *ten1*-*31* at all tested temperatures. Again, this suggested that the *ten1*-*31* mutation may affect transcription elongation, based on the fact that reducing the pools of UTP and GTP by 6-AU can compensate for defects in transcription elongation (Mason and Struhl 2005; Hartzog and Fu 2013). Accordingly, mutations in *RPB1* or in transcription factors that reduce RNA Pol II elongation rate are resistant to 6-AU or MPA (García et al. 2012; Braberg et al. 2013). We also used additional drugs known to confer hypersensitivity to transcription elongation mutants, such as 15 mM caffeine (Yao et al. 2000) and formamide at 2% (Prelich and Winston 1993). The *ten1*-*31* mutant was not sensitive to caffeine (**data not shown**) but, interestingly, was clearly hypersensitive to 2% formamide (**Fig. S4B**).

*Analysis of RNA pol II binding by ChIP PCR in* stn1-154

*stn1-154*, the more severe one of our collection of *stn1* mutants, only exhibited a slight but visible defect in RNA pol II binding at the 3’-end regions of the *PMA1* *and YEF3* long genes (**Fig. S5A, C**), but not in the case of the short *PGK1* gene (**Fig. S5A**). Therefore, Stn1 might contribute to the transcription functions of Ten1 studied here, as expected from its physical association with it (Grandin et al. 2001).

**Supplemental figure legends**

**Figure S1**. (***A-C***) *BUR1* and *CAK1* genetically interact with *TEN1*, but not with *STN1* or *CDC13*. (***A***) Genetic interactions between *TEN1* and *BUR1*, as well as between *TEN1* and *CAK1*, were observed as synthetic growth defects between the temperature-sensitive corresponding mutants, *ten1*-*31*, *bur1*-*80* and *cak1*-*23*. Growth of the *ten1*-*31 bur1*-*80* and *ten1*-*31 cak1*-*23* double mutants was strongly impaired when compared to that of each corresponding single mutant, an effect best seen at 32 and 34°C for *ten1*-*31 bur1*-*80* and already visible at 24°C for *ten1*-*31 cak1*-*23*. (***B***) The *ten1*-*6*, *ten1*-*16*, *ten1*-*3* and *ten1*-*100* mutants also exhibited synthetic growth defects when combined with the *bur1*-*80* mutation, effects that were best seen at 32°C, but that somewhat differed among *ten1* mutants because of the different degrees of severity of these mutants. Note that *ten1*-*6* and *ten1*-*3* are not temperature-sensitive, but yet genetically interact with *bur1*-*80*. (***C***) **Top panel:** Absence of genetic interactions between *CDC13* and *CAK1* or *BUR1* was observed after comparing growth of the *cdc13*-*1* *cak1*-*23* and *cdc13*-*1* *bur1*-*80* double mutants at different temperatures with that of the corresponding single mutants, as indicated. **Middle panel:** Absence of genetic interactions between *STN1* and *BUR1* was observed after comparing growth of the *stn1*-*13* *bur1*-*80* double mutant at different temperatures with that of the corresponding single mutants, as indicated. **Bottom panel:** Additional temperature-sensitive *stn1* mutants, *stn1*-*138*, *stn1*-*101* and *stn1*-*154*, also failed, like *stn1*-*13* above, to exhibit clear synthetic growth defects in combination with the temperature-sensitive *bur1*-*80* mutant, as indicated. (***D***) Directing expression of the *ten1*-*33* mutant allele under the control of *STN1*’s natural promoter instead of *TEN1*’s own promoter did not prevent the synthetic interactions between *ten1*-*33* and *bur1*-*80* from taking place.

**Figure S2**. *TEN1* exhibits genetic interactions with *KIN28*, but not with *SRB10*, nor mutants known to affect TERRA stability. (***A***) Schematic representation of the six CAK-controlled CDKs. (***B, C***) The *ten1*-*31* mutant did not exhibit synthetic interactions with a null mutation in the CDK-coding *SRB10* gene (***B***, only the most relevant temperature, 34°C, is shown here), while it did exhibit a synthetic interaction with a temperature-sensitive mutation in another CDK-coding gene, *KIN28* (***C***), best visible here at 29°C.

**Figure S3.** (***A***) Extension of data shown in Figure 1D. *TEN1* geneticaly interacts with *SPT4* and *SPT5*, as well as with *FCP1* and *RPB1*, as strong synthetic interactions between the temperature-sensitive *ten1*-*31* mutation and the temperature-sensitive *spt5*-*194*, *fcp1*-*1*, *rpo21*-*1* (a mutation in *RPB1*) mutations, as well as the *spt4* null mutation (*spt4Δ*), are observed. *TEN1* also exhibited strong genetic interactions with *CDC73*, coding for a component of the PAF1 complex. (***B, C***) Ten1 does not appear to function in TERRA regulation. Null mutations in *THP2* or *MFT1*, coding for two components of the TERRA-regulating THO complex did not synthetically interact with *ten1*-*31 (****B****)* nor were null mutations in *RNH1* and *RNH201*, coding for the two RNase H nucleases responsible for the removal of R-loops accumulating in mutants of the THO complex (***C***).

**Figure S4**. (***A***) **Top two panels** show that the *ten1*-*31* mutant was resistant to 6-AU, similarly to the wild type (*wt*), but unlike the *spt4*Δ mutant. Interestingly, the *ten1*-*31* mutation suppressed *spt4*Δ sensitivity to 6-AU, although it clearly increased the growth defect of *spt4*Δ at 29°C. **Bottom panel** illustrates RT-PCR showing that *ten1*-*31* cells constitutively express the *IMD2* gene, independently of 6-AU treatment (50 μg/ml for 1 h), and this was confirmed by RNA-seq. *ACT1*, whose expression does not depend on the presence of 6-AU, was used as a control, and 18S rRNA as an independent control of RNA Pol I transcription. PCR products were run in a 1.5% agarose gel stained with EtBr. (***C***) *ten1-31* mutant cells are sensitive to formamide (FA) under conditions indicated in the figure. Cells were grown at different temperatures in order to take into account their respective degree of temperature sensitivity.

**Figure S5.** (***A***) ChIPs to analyze Rpb3 association to several constitutively transcribed genes, *PMA1* and *PGK1* in wt, *ten1*-*31*, *stn1*-*154* and *spt5*-*194*. (***B***) The *ten1-31* mutation does not affect Rpb1 nor Rpb3 total protein levels, but causes a slight increase in Ser2P levels. Whole cell extracts (WCE) were prepared from wild-type (*wt*) and *ten1-31* strains and analyzed by western blotting using the following antibodies: anti-Rpb1 (8W16G), anti-Rpb3, anti-Ser2P (3E10), and anti-Pgk1 as a control for total protein. (***C***) Rpb3 occupancy at *PGK1*, *PMA1* and *YEF3* genes in wt, *ten1*-*31* and *stn1*-*154* cells.

**Figure S6.** Relative levels of Ser2P/Rpb1 are increased in *ten1-31* cells nearby the promoters in most of the tested genes, at the 5’ coding regions. Ser2P/Rpb1 ratio was calculated using values obtained from the Rpb1 and Ser2P ChIP-qPCR assays represented in Figure 2 for all five genes tested, *PMA1, YEF3, PGK1* (***A***), *FMP27 and GAL1* (***B***).

**Figure S7**. Gene length-dependent Rpb1-Ser2P levels are altered in *ten1* mutant cells. (***A***) **Left panel:** enrichment of Rpb1-Ser2P over total RNA pol II along protein-coding gene regions in wt and *ten1-31* cells for short genes (0-1 Kb, n = 1182). **Right panel:** Rpb1-Ser2P levels in *ten1-31*/wt for short genes. (***B***) Same as (A) for medium-sized genes (1-2 Kb, n = 2319). (***C***) Same as (A) and (B) for long genes (> 2 Kb, n = 1639). Standard deviations are represented as translucent areas around the solid traces.

**Figure S8**. ChIP-seq signal in selected genomic loci. Genome browser maps around *PMA1*, *YEF3* and *PGK1*. **Blue track:** differential Rpb1 binding in *ten1-31* mutant compared to wt. **Gold track:** same for the Rpb1-Ser2P ChIP signal. **Green track:** enrichment of Rpb1-Ser2P signal over total RNA pol II (Rpb1) signal in wt. **Orange track:** same in *ten1-31*. The values in top left corners correspond to the log_2_ of the number of reads ratio.

**Figure S9.** Correlation analysis of RNA-seq replicate datasets. The expression levels of the protein-coding genes in the different samples were compared against each other and plotted as a correlation matrix. Spearman correlation values are displayed for each pairwise comparison. TEN1 and WT refer to the *ten1-31* or wt strains, respectively. Warmer colours correspond to a higher density of overlaying dots (genes).

**Figure S10**. Cdc13-Myc_13_ and Ten1-Myc_13_, but not Xrs2-Myc_13_, associate with Spt5. Xrs2, which has a size similar to that of Cdc13, is part of the so-called MRX (Mre11-Rad50-Xrs2) complex implicated in NHEJ repair. Co-IP assays were performed in the Cdc13-Myc_13_,Ten1-Myc_13_ and Xrs2-Myc_13_ Myc-tagged strains. Input and IPs were analyzed by western blotting with antibodies (anti-Myc or anti-Spt5) to the indicated proteins.

**Figure S11.** ChIPs to analyze RNA Pol I association to the rDNA. The graph shows Rpa190 occupancy at the promoter of the 35S rDNA and within the 25S and 18S. ChIP analyses, quantification and graph representation were done as in the other cases illustrated above. Thus, the values obtained for the IPed PCR products were compared to those of the total input, but in this particular case, the values from each PCR product from the transcribed regions were normalized to the 5S rDNA.

**Figure S12.** Two-hybrid interactions between Pol1 and Spt5, as well as between Cdc13 and Stn1 or Pol1. (***A***) Y190 strains simultaneously expressing pAS2-POL1-first 1146 nt and pACT2-SPT5-first 630 nt (**top row**) or pACT2 alone (**botton row**) were positive for β-galactosidase activity in the X-gal assay, (***B***) like cells coexpressing pAS2-CDC13 and pACT2-STN1 or pACT2-POL1-1146, unlike those expressing pACT2 alone, used here as negative control. Patches of cells replica-plated on nitrocellulose membrane were incubated for 2 h at 30°C and then photographed.

**Figure S13.** *TOP1* and *TOP2* genetically interact with *TEN1*, but only when both *TOP1* and *TOP2* have been mutated. The *top1*Δ and temperature-sensitive *top2*-*1* mutants were used. Growth of the *ten1*-*31 top2*-*1* *top1* Δ triple mutant was clearly impaired when compared to that of each corresponding single or double mutants.

**Figure S14.** Correlation analysis of ChIP-seq replicate datasets. Sequencing datasets were compared against each other after alignment against the reference genome. Spearman correlation values are displayed for each pairwise comparison. TEN1 and WT refer to the *ten1-31* or wt strains, respectively. Ser2 and Rpb1 refer to the IPs against the Ser2 phosphorylated form of Rpb1 or the total Rpb1, respectively.

**STAR METHODS**

**Materials and methods**

*Yeast strains and media*

*Saccharomyces cerevisiae* yeast strains used in this study were derivatives of BF264-15Daub (*ade1* *his2* *leu2*-*3*,*112* *trp1*-*1a* *ura3Dns*), and as well as yeast culturing, have been described previously (Grandin et al. 1997). All strains were made isogenic by back crossing at least five times against our genetic background.

The *ten1*-*16* (F154I) and *ten1*-*31* (E58K, L76P, E91V and V115A) mutants have been previously described (Grandin et al. 2001). The *ten1*-*33* harbors the K40E, I44M, K55E and L76P mutations. All three *ten1* mutants are temperature-sensitive at different levels, exhibiting more or less growth impairement at 36°C. None of these three mutants exhibit a tight arrest, and all of them exhibit morphological defects at all temperatures comprised between 24 and 36°C, at various levels depending on the allele concerned. In all *ten1* mutants, the ORF was under the control of its natural promoter, expressed from a centromeric plasmid of the YCplac series (Gietz and Sugino 1988) in strains in which *TEN1* had been completely deleted (Grandin et al. 2001). It is important to note that *ten1*-*16*, *ten1*-*31* and *ten1*-*33* were totally inviable when present at single copy following expression from an integrative YIplac vector, as reported before (Grandin et al. 2001), and had, therefore, to be expressed from a YCplac vector (Gietz and Sugino 1988). Such CEN vectors are known to express genes under their control at 2-4 copies (Rose et al. 1987). This was not the case for *ten1*-*3*, *ten1*-*6* and *ten1*-*13*, which are not temperature-sensitive mutants and were isolated on the basis of conferring elongated telomeres and can be expressed under viable conditions from an integrative vector (Grandin et al. 2001).

The viability of cells previously grown in liquid was determined by performing and analyzing the so-called “drop tests” or “spot assays”. To do this, cells from exponential growth cultures were counted with a hematocytometer and the cultures were then serially diluted by 1/5th or 1/10th and spotted onto YEPD (or selective medium) plates and incubated at the desired temperatures for 2–3 days before being photographed. In some cases, cells were just re-streaked onto YEPD plates and growth evaluated by visualizing the numbers and sizes of the growing colonies.

*Genetic screen to find extragenic mutations enhancing the* ten1 *phenotype*

For strain mutagenesis, *ten1* mutant strains were grown overnight, almost to saturation, in liquid YEPD medium at 24°C, before being diluted 1/5,000^th^ to 1/7,500^th^ in 1 ml H*_2_*O, from which 200 μl were plated out onto solid YEPD plates, which were then exposed to UV light (254 nm wave length) at a distance of 10-15 cm for 6-10 s. Plates of UV-mutagenized yeast strains were then incubated at 24°C for ~ 50 h before being replicated on YEPD plates at 36°C. After 1.5-2.5 days, the 24°C and 36°C replicas were compared between them; colonies failing to grow well at 36°C were selected and their 24°C counterpart repatched at 24°C before being re-tested for growth at 36°C.

*Co-immunoprecipitation and western blot analysis*

A strain expressing Ten1-Myc_13_ or Cdc13-Myc_13_ , or Xrs2-Myc_13_ strain for a negative control, were grown in 200 ml of YEPD to an OD600 of 1.5, harvested, washed with water, and suspended in 2.5 ml of lysis buffer (20 mM HEPES pH 7.6, 200 mM potassium acetate, 1 mM EDTA pH 8.0, glycerol 10%) containing protease and phosphatase inhibitors. The cell suspension was flash frozen in liquid nitrogen, and then ground in a chilled mortar to a fine powder. Afterwards, the cell lysate was thawed and centrifuged at 13,200 rpm for 20 min. The supernatant was collected and total protein concentration was estimated measuring absorbance at 280 nm in a nanodrop. Prior to immunoprecipitation, the extracts were precleared by incubation with either protein G agarose (Santa Cruz) or protein A sepharose (GE healthcare) for 1 h to eliminate as much as possible unspecific binding. Then, they were centrifugated and the supernatants used for either anti-Myc or anti-Spt5 immunoprecipitations. For that, the volume of each cell extract containing 20 μg of protein was incubated with 1 μl of anti-Myc (Roche) or 2.5 μl of anti-Spt5 (Santa Cruz) for 1 h at 4ºC, and afterwards 20 μl of protein A sepharose or protein G agarose slurry, respectively, were added and the incubation extended to overnight at 4°C. The IPs were extensively washed with lysis buffer and beads were suspended in SDS-PAGE sample buffer. Thereafter they were incubated at 65°C for 20 min and supernatants were loaded onto a SDS-PAGE gel and analyzed by western blot with the corresponding antibodies. For Ten1-Myc_13_/Hmo1-HA_2_ co-IP assays, we proceeded as above. In that case, 20 μg of whole cell extracts from Ten1-Myc_13_, Hmo1-HA_2_ and Ten1-Myc_13_ Hmo1-HA_2_ strains were IPed with 2 μl of anti-Myc, and then assayed by western blot using anti-HA and anti-Myc antibodies.

*Two-hybrid experiments*

The two-hybrid "kit" was kindly provided by Stephen J. Elledge. Experiments of protein-protein interactions using the two-hybrid system were performed as described previously (Fields and Song 1989; Durfee et al. 1993; see also Grandin et al. 1997). Genes of interest were cloned in-frame with the *GAL4* activation domain (nucleotides 764-885) in pACT2 or in-frame with the *GAL4* DNA-binding domain (nucleotides 1-147) in pAS2. Both types of constructs were transformed into the Y190 strains, which were then tested for β-galactosidase activity, toghether with the appropriate controls, as described previously (Durfee et al. 1993) using X-gal (5-bromo-4-chloro-3-indolyl-p-D-galactopyranoside, from Sigma).

*RNA isolation and RT-PCR*

Total RNA was extracted as described (Garavís et al. 2017) and RT-PCR was performed using the iScript RT reagent Kit (Bio-Rad), following the manufacturer’s instructions. PCR reactions were performed in triplicate with at least three independent cDNA samples.

*RNA-seq*

*1- Library preparation, sequencing and quality control*

Yeast total RNA was extracted from *ten1*-*31* mutant and wild-type (wt) cells after shifting to restrictive temperature, 34°C, for 2 hr. RNA from three biological replicates of the experiment was prepared independently for each condition (total of 6 samples) and subjected to rRNA depletion. The rRNA depleted fraction was then used to construct libraries that were sequenced in a HiSeq system (Ilumina) to an output of 120 (1 x 50 nt) million strand-specific reads. Raw reads were analyzed with FastQC to make sure each sequenced sample met the adequate quality standards.

*2- Adapter clipping, quality trimming and filtering*

Trimmomatic (Bolger et al. 2014) was used to clip, filter and remove certain portions of reads or even entire reads. The Illuminaclip function was used to remove Illumina adapters. The SLIDINGWINDOW function was applied to filter out reads shorter than 20 bases and the MINLEN function was used to discard reads with an average quality of less than 28. A second round of inspection with FastQC was carried out after the filtering steps to ensure that the filters were applied properly.

*3- Genome alignment and expression matrix generation*

Filtered reads were mapped to the yeast genome with TopHat2 (Kim et al. 2013), using the R64 (sacCer3) genome as reference. 94.7% of total reads were successfully aligned to the reference genome. A table with information about *per sample* total number of reads, trimmed and quality-filtered reads and overall alignment rates is provided as **Table S1**. BAM alignment files were visually inspected with the genome browser IGV ([www.broadinstitute.org/igv/](http://www.broadinstitute.org/igv/)). Normalized coverage tracks for genome browser visualization were generated with the functions bamCoverage and bamCompare from the deepTools2 suite (Ramírez et al. 2016). Coverage tracks from individual samples are expressed as reads per kilobase per million mapped reads (RPKM), whereas comparison tracks are expressed as the log_2_ of the number of reads ratio. Raw read counts were extracted with the featureCounts tool from the R package Rsubread (Liao at al. 2014), using the yeast genome annotation as a reference.

*4- Similarity of biological replicates*

The similarity of biological replicates was assessed with correlation analysis after log_2_ transformation of the raw read expression matrix. The function heatpairs from the R package LSD (<https://CRAN.R-project.org/package=LSD)> was used to generate a matrix of pairwise correlation scatterplots. Spearman correlation coefficients are reported by default for each comparison (**Fig. S9**). The raw expression matrix was used as input for differential expression analysis.

*5- Differential expression analysis and GSEA*

The list of differentially expressed genes between samples was obtained by using DESeq2 after applying a 0.3 non-differential contig count quantile threshold and a 0.05 p-value threshold for the FDR control test. Changes in expression between *ten1*-*31* and wt samples are expressed as log_2_ Fold Change (mutant/wt). The list of genes with log_2_ FC values as calculated by DESeq2 was ordered from highest to lowest and used as input to test for GO category enrichment, as implemented in GOrilla (Eden et al. 2009).

*Chromatin immunoprecipitation (ChIP) and ChIP-seq*

*1- ChIP by qPCR*

Chromatin purification, immunoprecipitation, quantitative real-time PCR (qPCR) amplification and data analysis were performed as described (García et al. 2010; Allepuz-Fuster et al. 2014; Garavís et al. 2017). Briefly, cells were grown overnight to saturation at 25ºC, then diluted to an OD_600_ of 0.2 approximately, and let them grow at 34 ºC for almost 5 h until they reached an OD_600_ of 0.5-0.6. Immunoprecipitated and purified chromatin was subjected to quantitative real-time PCR using the CFX96 Detection System (Bio-Rad Laboratories, Inc.) and SYBR® Premix Ex Taq ™ (Takara Bio), following the manufacturer’s instructions. Real-time PCR reactions were performed in duplicate from at least three independent ChIPs. Quantitative analysis was performed with the CFX96 Manager Software (version 3.1, Bio-Rad). The values obtained for the IPed PCR products were compared to those of the total input, and the ratio of the values from each PCR product from transcribed genes to a non-transcribed region of chromosome VII was calculated. Numbers on the y-axis of graphs are detailed in the corresponding figure legend.

*2- ChIP-seq*

*2-1- Library preparation, sequencing and quality control*

For ChIP-seq experiments, anti-mouse or anti-rat IgG dynabeads (Invitrogen) were used to immunoprecipitate Rpb1 or Rpb1-Ser2P, together with 8WG16 and 3E10 antibodies, respectively. Each experiment was carried out in biological duplicates for a total of 12 samples. Library DNA was prepared from immunoprecipitated DNA and its corresponding input DNA following the manufacturer's instructions, and was sequenced in a Illumina HiSeq 2500 system to an output of 313 (1 x 50) million reads. The quality metrics of the fastq sequencing datasets were obtained with FastQC and visually inspected.

*2-2- Filtering, trimming and adapter clipping*

These steps were carried out essentially as described for RNA-seq.

*2-3- Genome alignment and genome browser visualization*

High quality reads were mapped to the yeast genome with Bowtie2 (Langmead and Salzberg 2012), using the R64 (sacCer3) genome as reference. A table with alignment statistics is provided as **Table S2**. BAM alignment files were visually inspected with the genome browser IGV. Normalized coverage tracks for genome browser visualization were generated with the functions bamCoverage and bamCompare from the deepTools2 suite (Ramírez et al. 2016). Coverage tracks from individual samples are expressed as reads per kilobase per million mapped reads (RPKM), whereas comparison tracks are expressed as the log_2_ of the number of reads ratio.

*2-4- Reproducibility of the replicates and merging of datasets*

A matrix of read coverages for the entire genome was generated with all alignment files with the function multiBamSummary from deepTools2. Briefly, the function takes the genome annotation and splits it in 10 kilobase bins. For each bin, the number of reads found in each bam dataset is counted. A correlation matrix heatmap plot was generated with the plotCorrelation function of the same package. Correlation values are calculated with the Spearman method as default (**Fig. S14**). Based on the high correlation values of the biological replicates, the two datasets from each sample were merged into one for average metagene representations around genomic regions with SamTools (Li et al. 2009).

*2-5- Metagene analysis*

Normalized average density plots around genomic features were calculated with the ngs.plot software (Shen et al. 2014). The ChIP-seq mode and the statistical robustness parameter, which filters out 0.5% of genes with the most extreme expression values, were applied to all calculations.

*2-6- ChIP-exo of Hmo1*

Hmo1 ChIP-exo datasets were downloaded from the Sequence read archive (SRA) repository, with accession number SRP041518. Raw fastq files were processed following a pipeline similar to the one described above for our ChIP-seq datasets.

*Mass spectrometry analysis*

Anti-Myc immunoprecipitates from a Ten1-Myc_13_ (or Stn1-Myc_13_) strain or control isogenic untagged strain were loaded in a SDS-PAGE acrylamide concentrator gel and run for a very short time, after which, the sample-containing bands were cut out and digested with trypsin using and automatic digestion robot (Bruker Proteineer) under reducing (DTT) conditions. Carbamidomethylation of the samples was then performed using iodoacetamide. After digestion, samples were dried and dissolved in a final volume of 15 μl. A total volume of 5 μl of each sample was analyzed by liquid chromatography (short gradient, 100 min) coupled to a QTOF mass spectrometer (5600 Triple-TOF) along which the peptides were separated as a function of their hydrophobicity (using a C-18 reversed phase column). The eluted peptides were then fragmented on the 5600 Triple-TOF mass spectrometer, thereby obtaining a high number of MS and MSMS spectra of the precursors (peptides) present in the sample. Raw data were exported, processed and used to launch a search against the Uniprot *Saccharomyces cerevisiae* database using the MASCOT search engine. Peptide identifications with Mascot scores equal or greater than 20 were exported and used to establish the statistical significance of the identifications. False discovery rates (FDR) were applied at different levels: spectrum, peptide and protein, among which the protein level was considered for analysis. As generally accepted, a FDR value was considered to be significant when smaller than –or equal to- 1%.
