## Supplementary figures and images for "The telomeric Cdc13-Stn1-Ten1 complex regulates RNA polymerase II transcription"

### Supplementary file 2

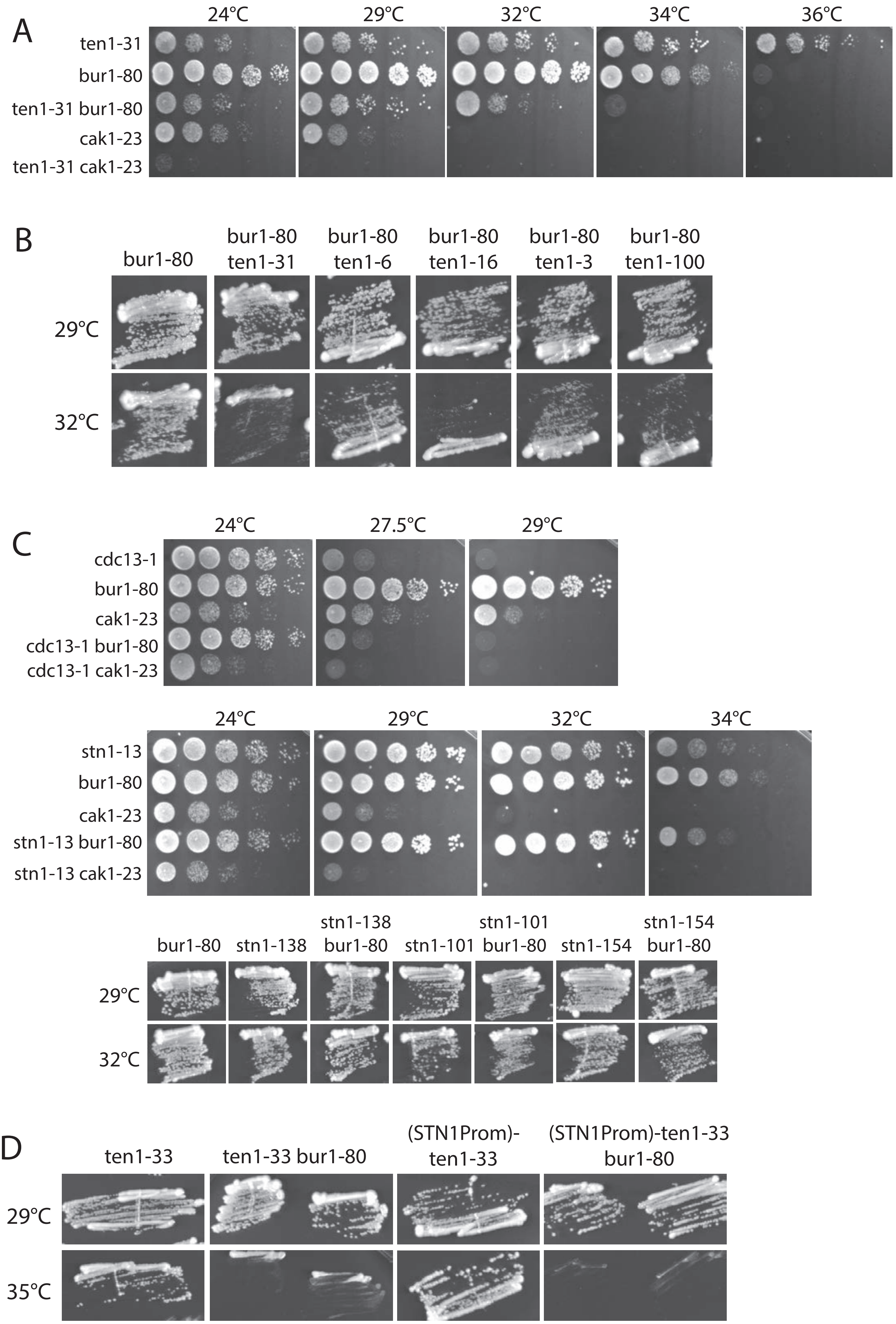

### Supplementary file 3

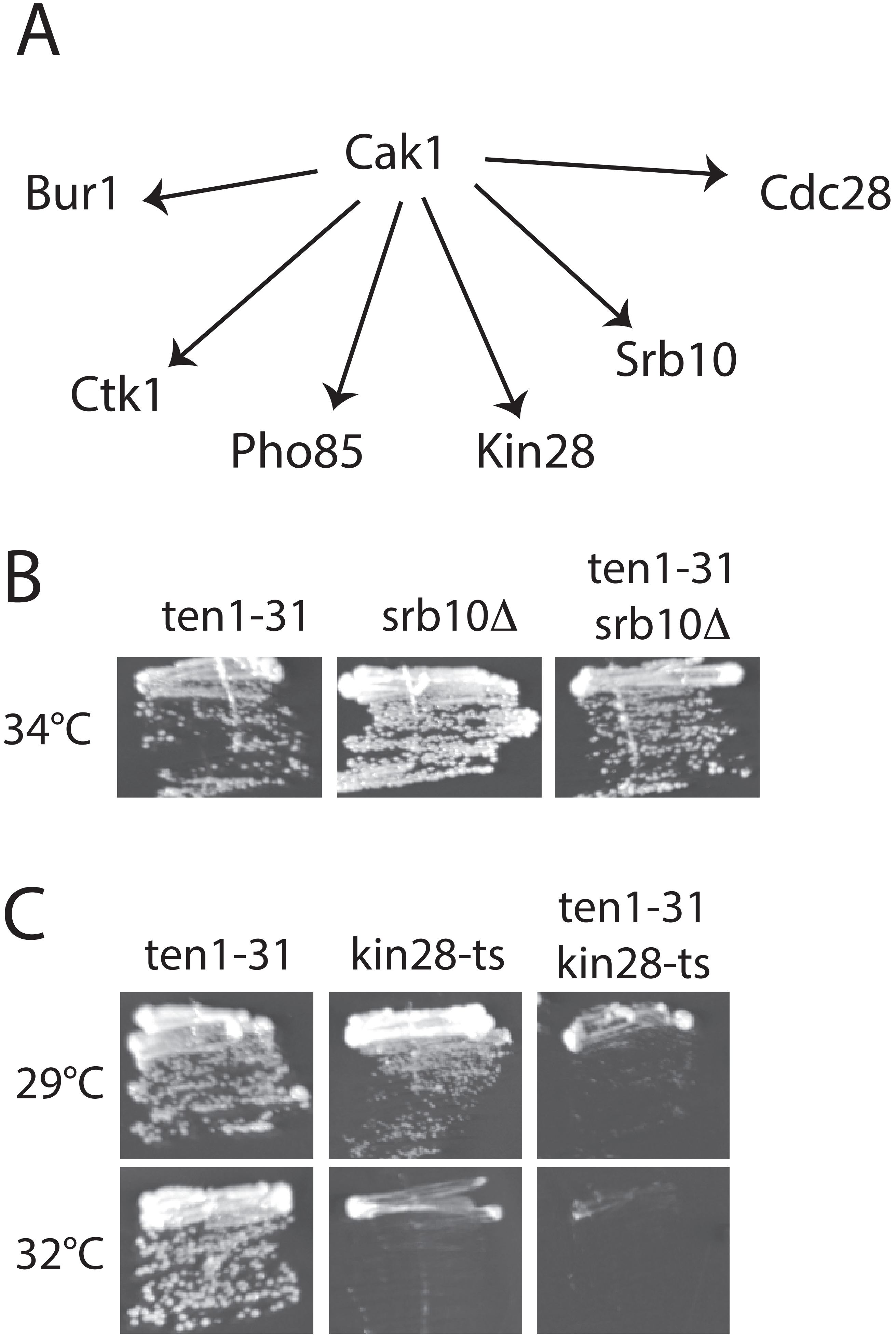

### Supplementary file 4

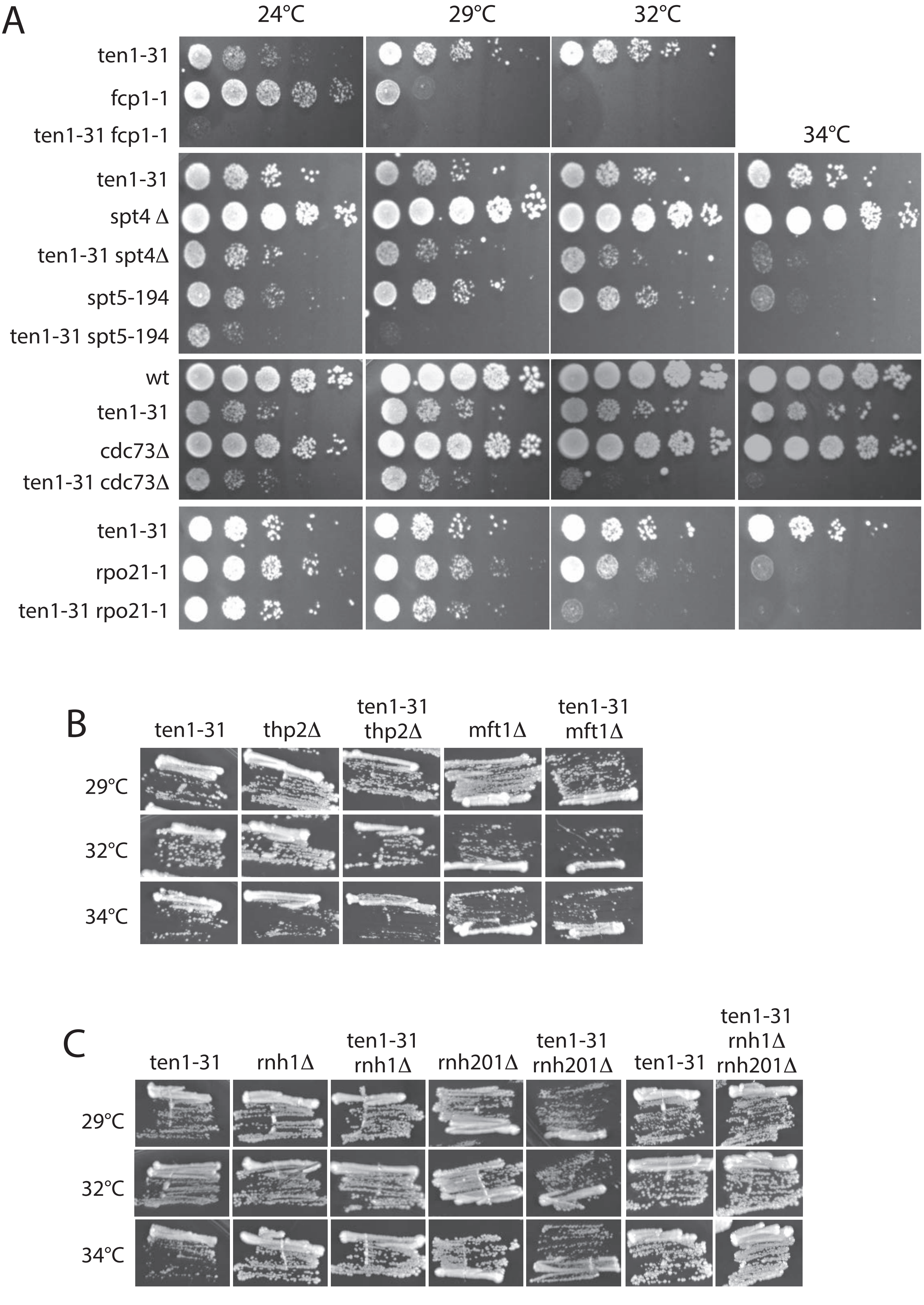

### Supplementary file 5

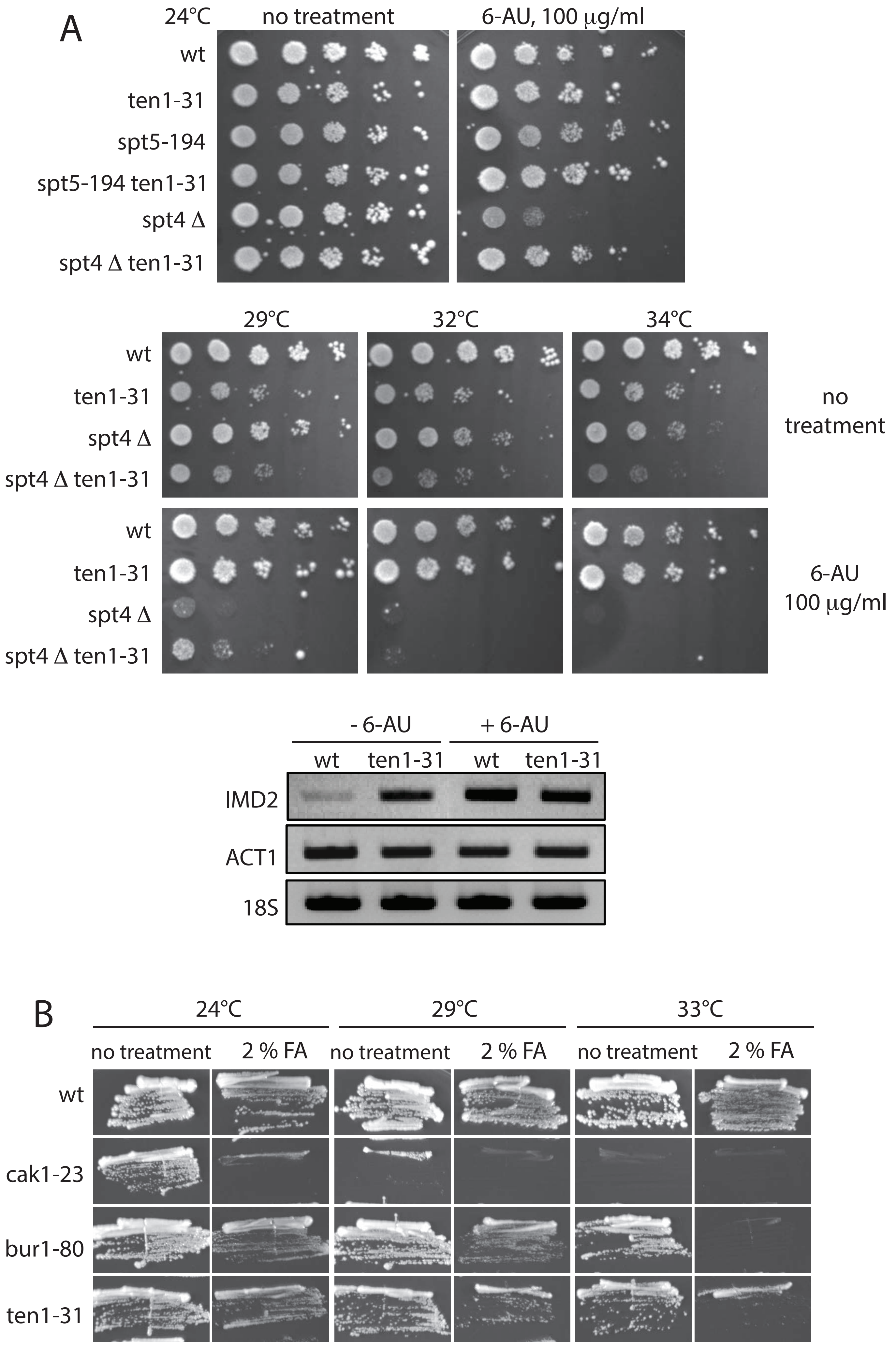

### Supplementary file 6

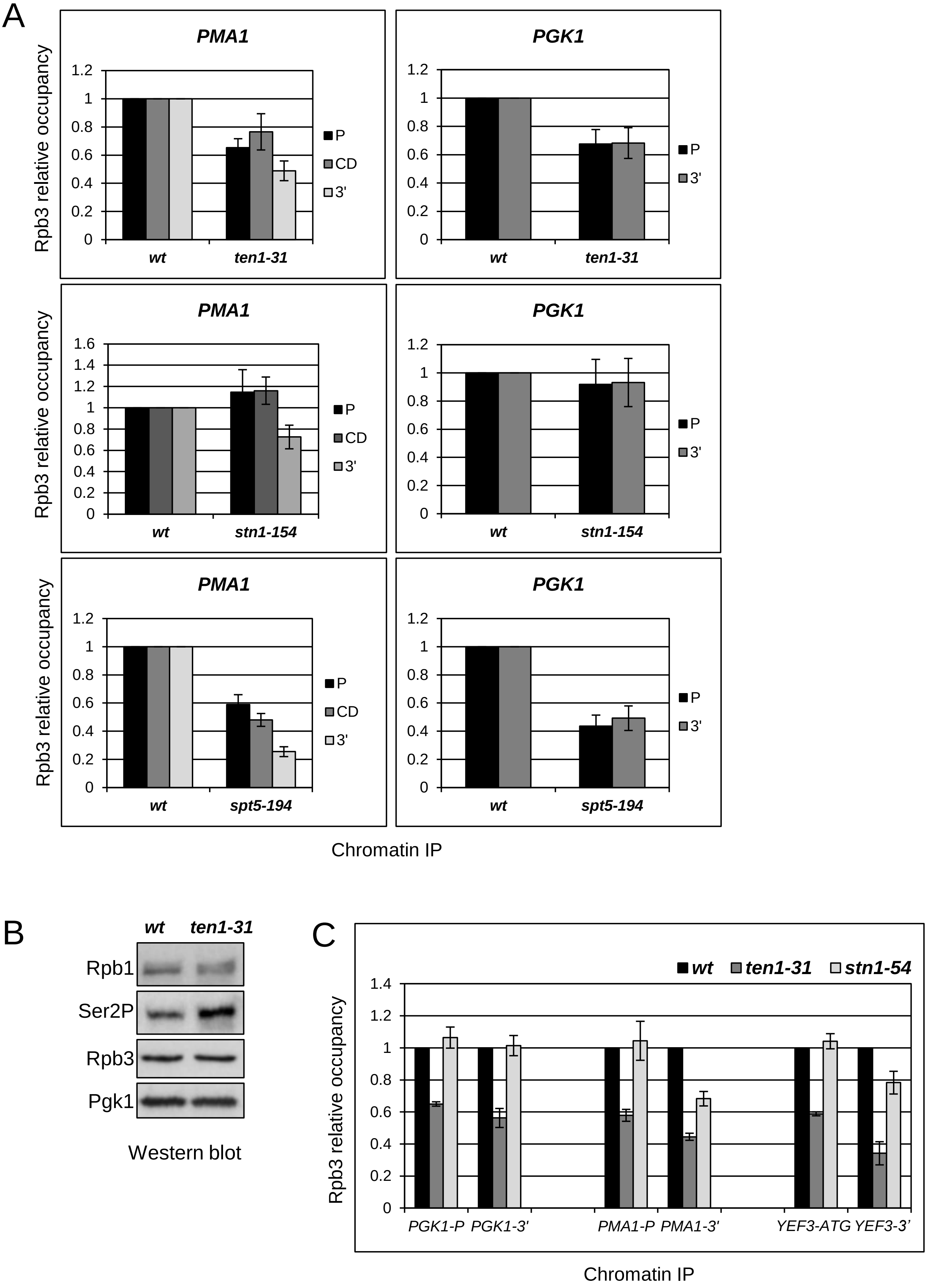

### Supplementary file 7

**Ser2P / Rpb1 ratio**

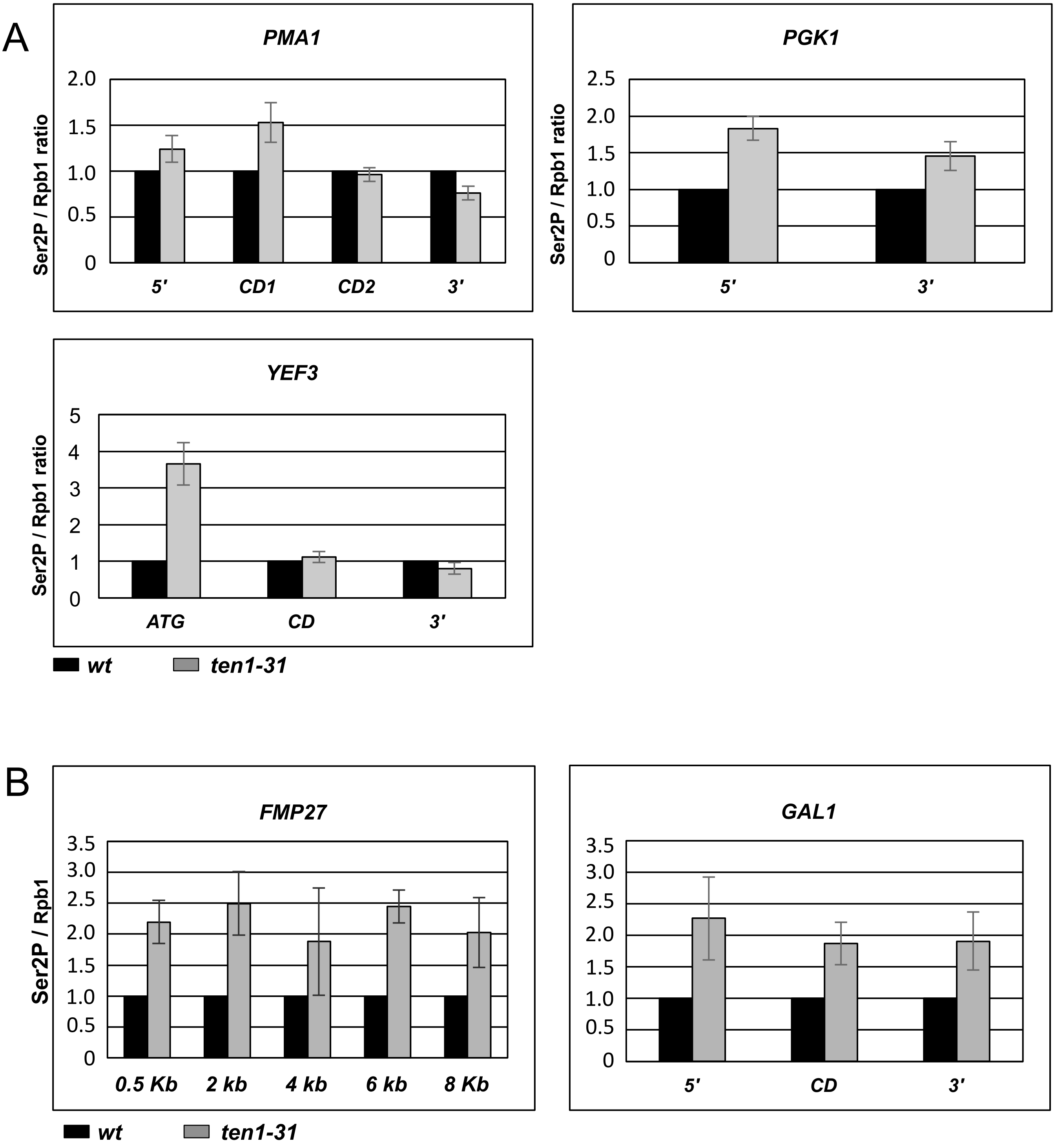


Figure S6

### Supplementary file 8

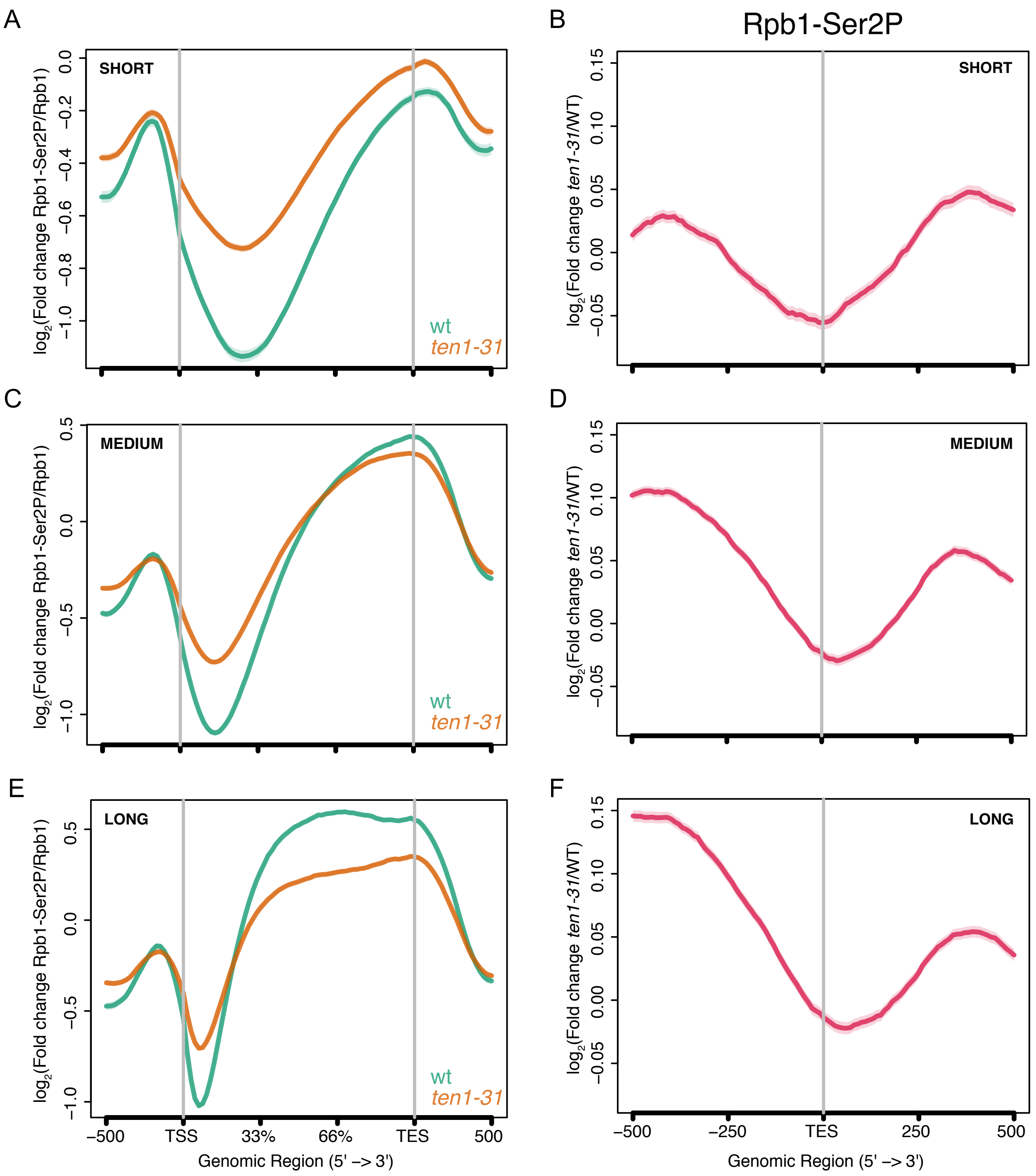

### Supplementary file 9

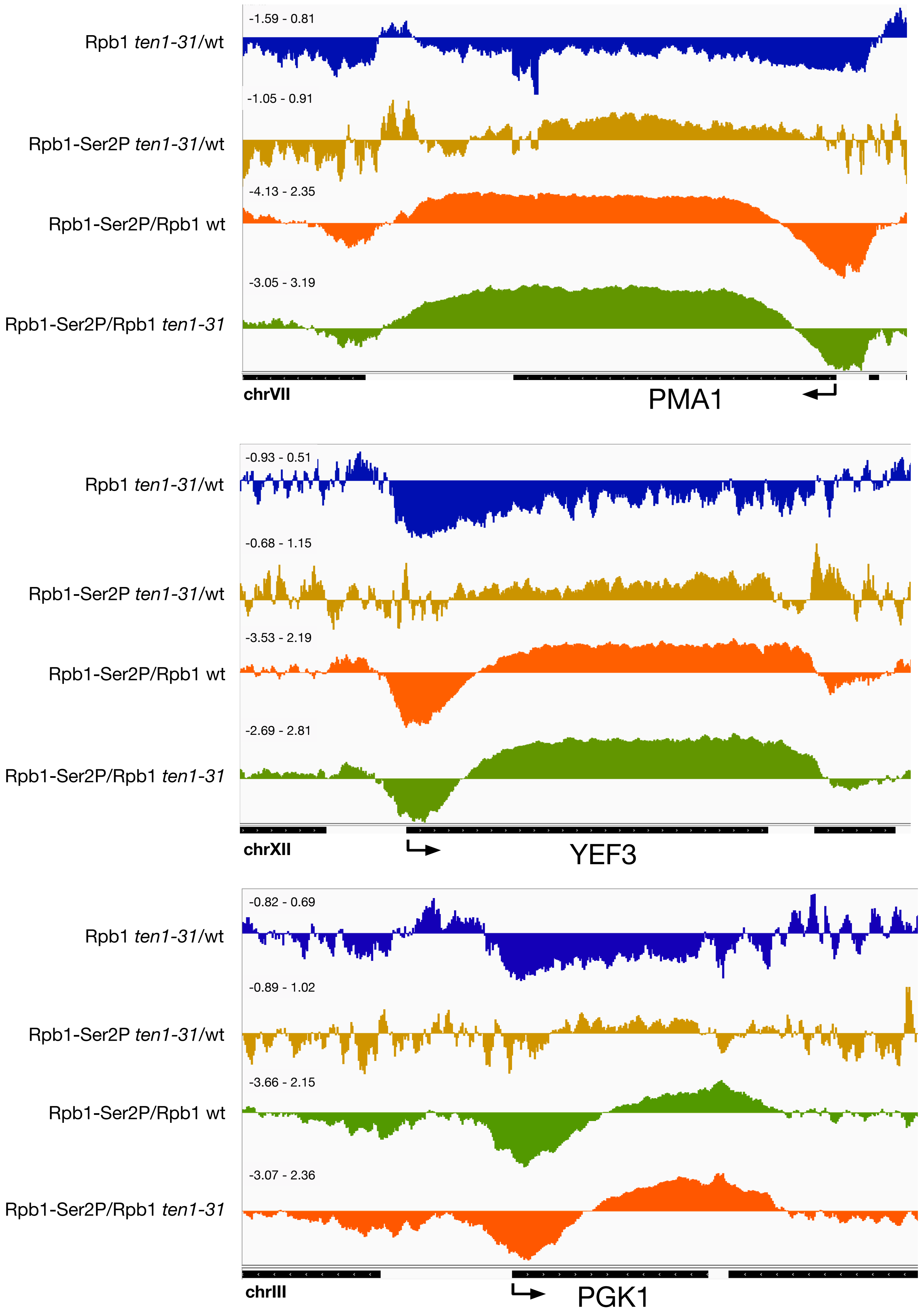

### Supplementary file 10

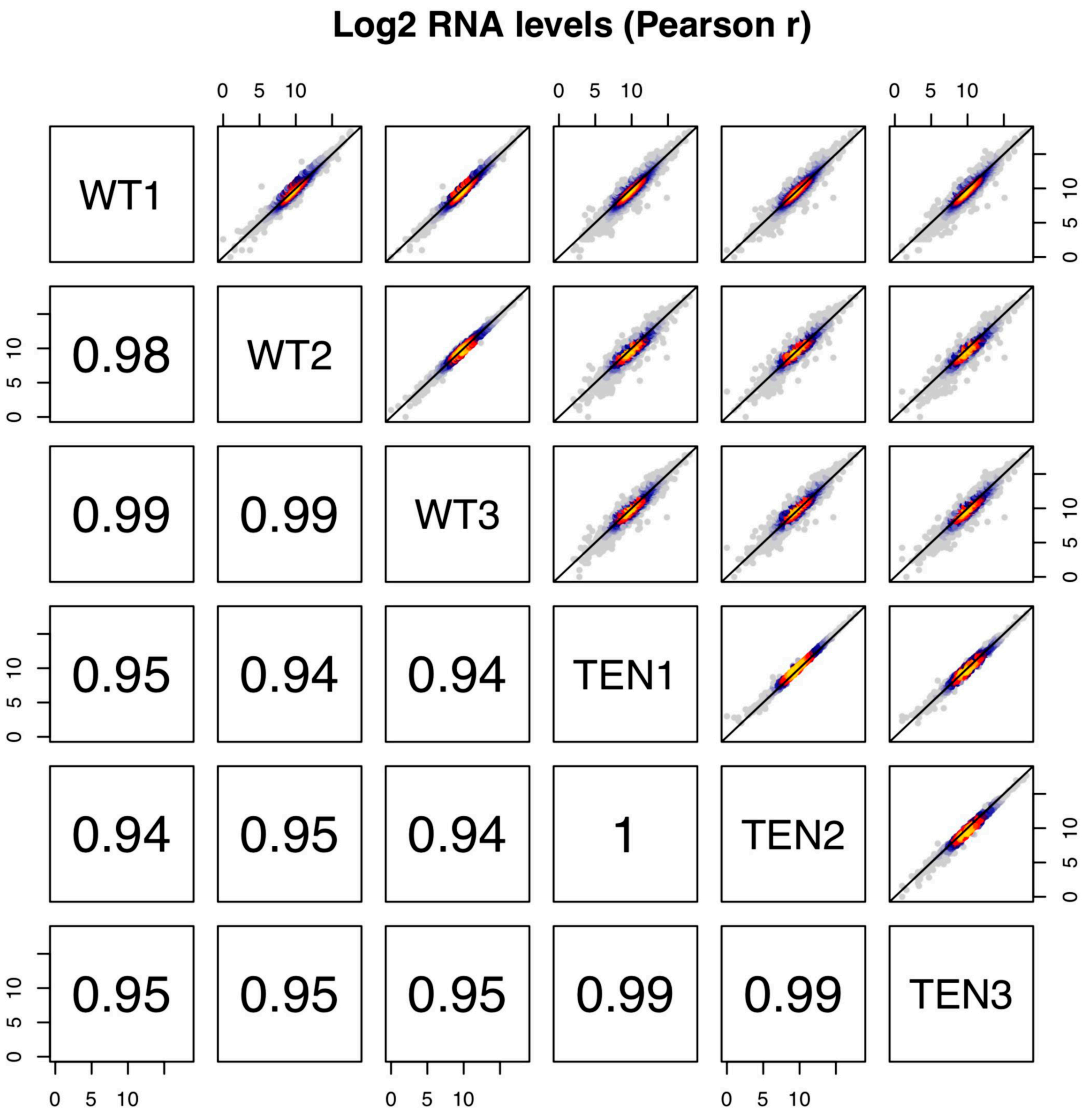

### Supplementary file 11

#### Slide 1
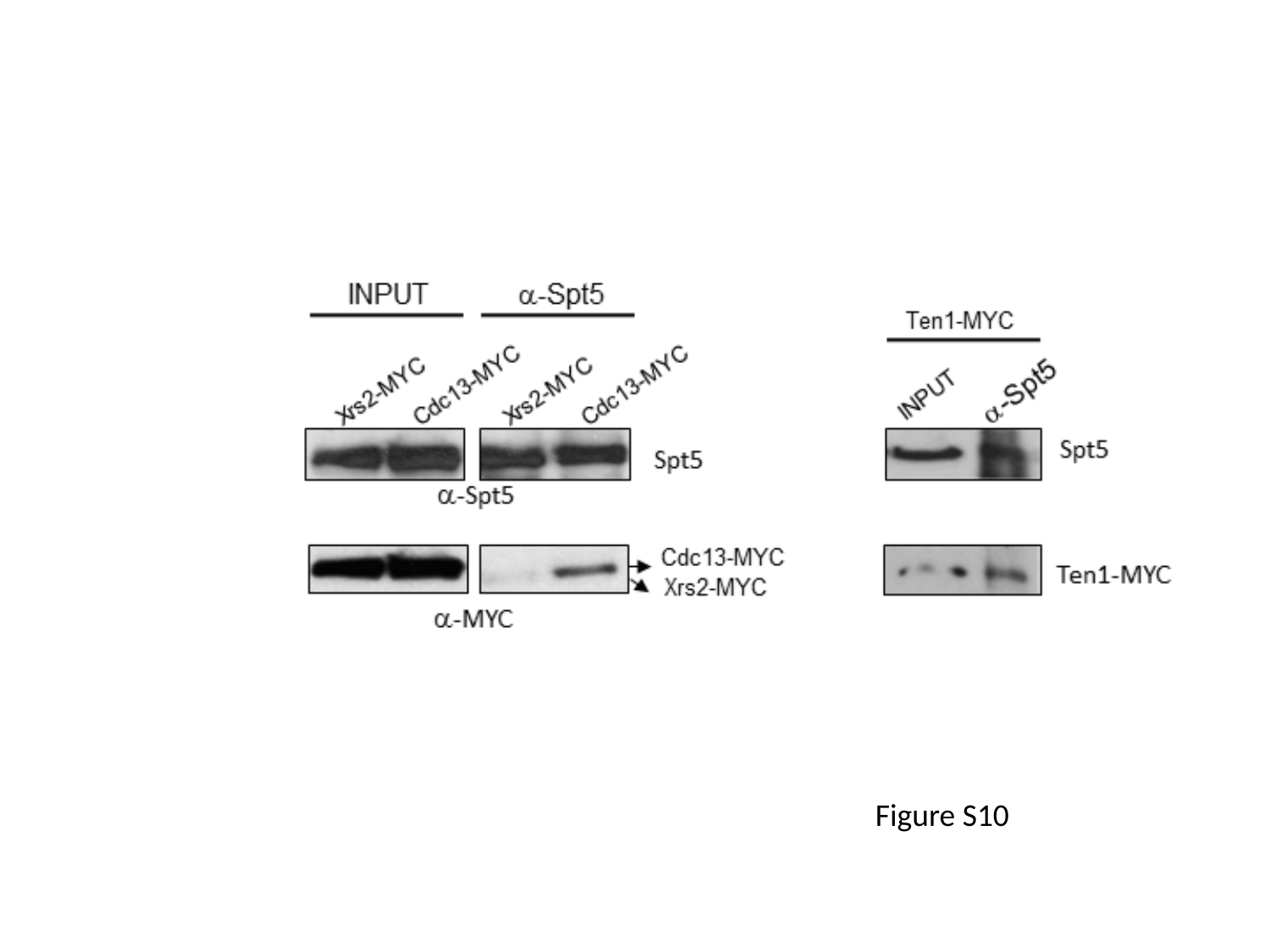

Figure S10

### Supplementary file 12

#### Slide 1
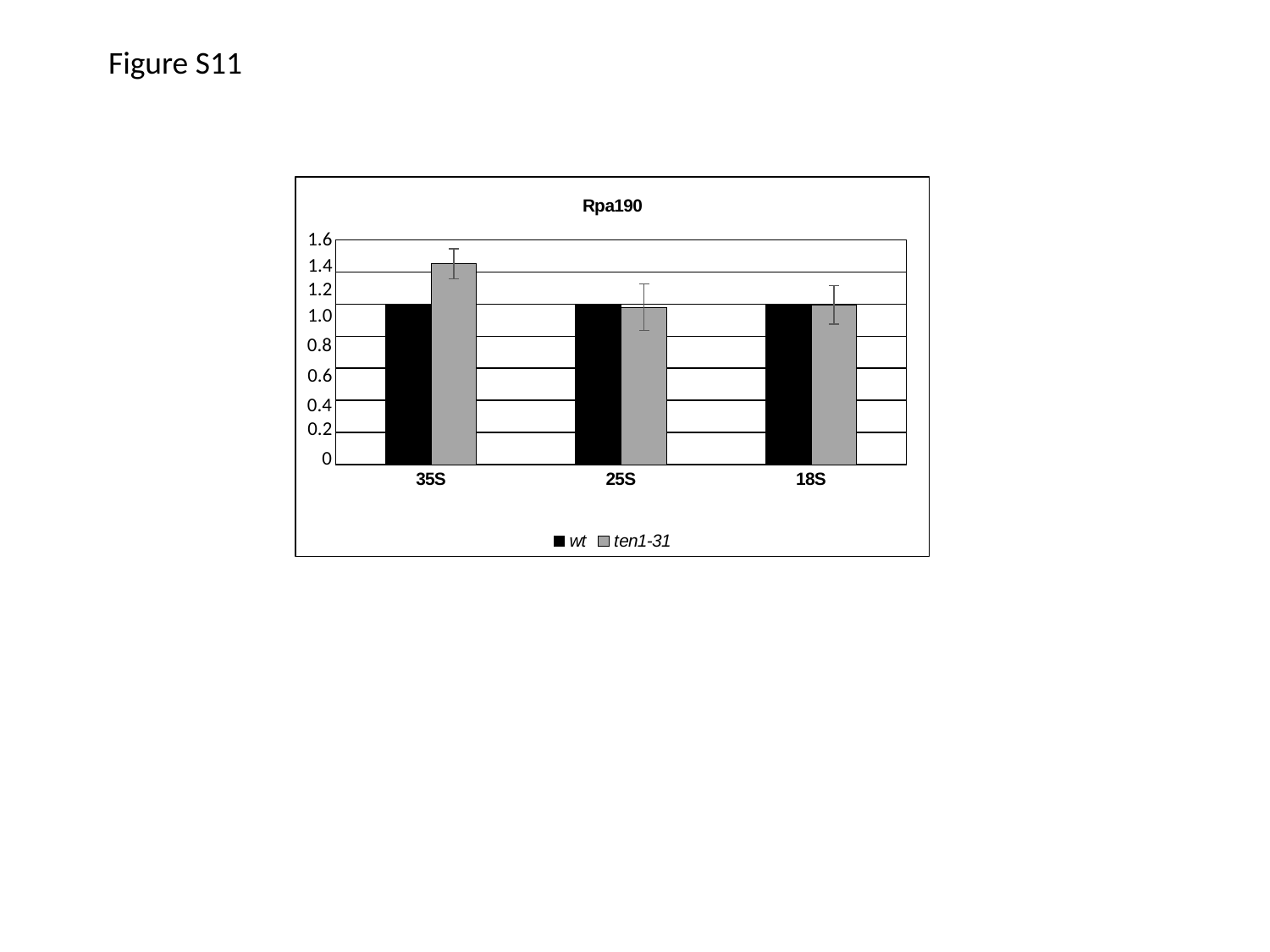

Figure S11
##### Chart: Rpa190
| Category | wt | ten1-31 |
|---|---|---|
| 35S | 1.0 | 1.25121621023681 |
| 25S | 1.0 | 0.980830332116576 |
| 18S | 1.0 | 0.994756947543699 |1.6
1.4
1.2
1.0
0.8
0.6
0.4
0.2
0

### Supplementary file 14

#### Slide 1
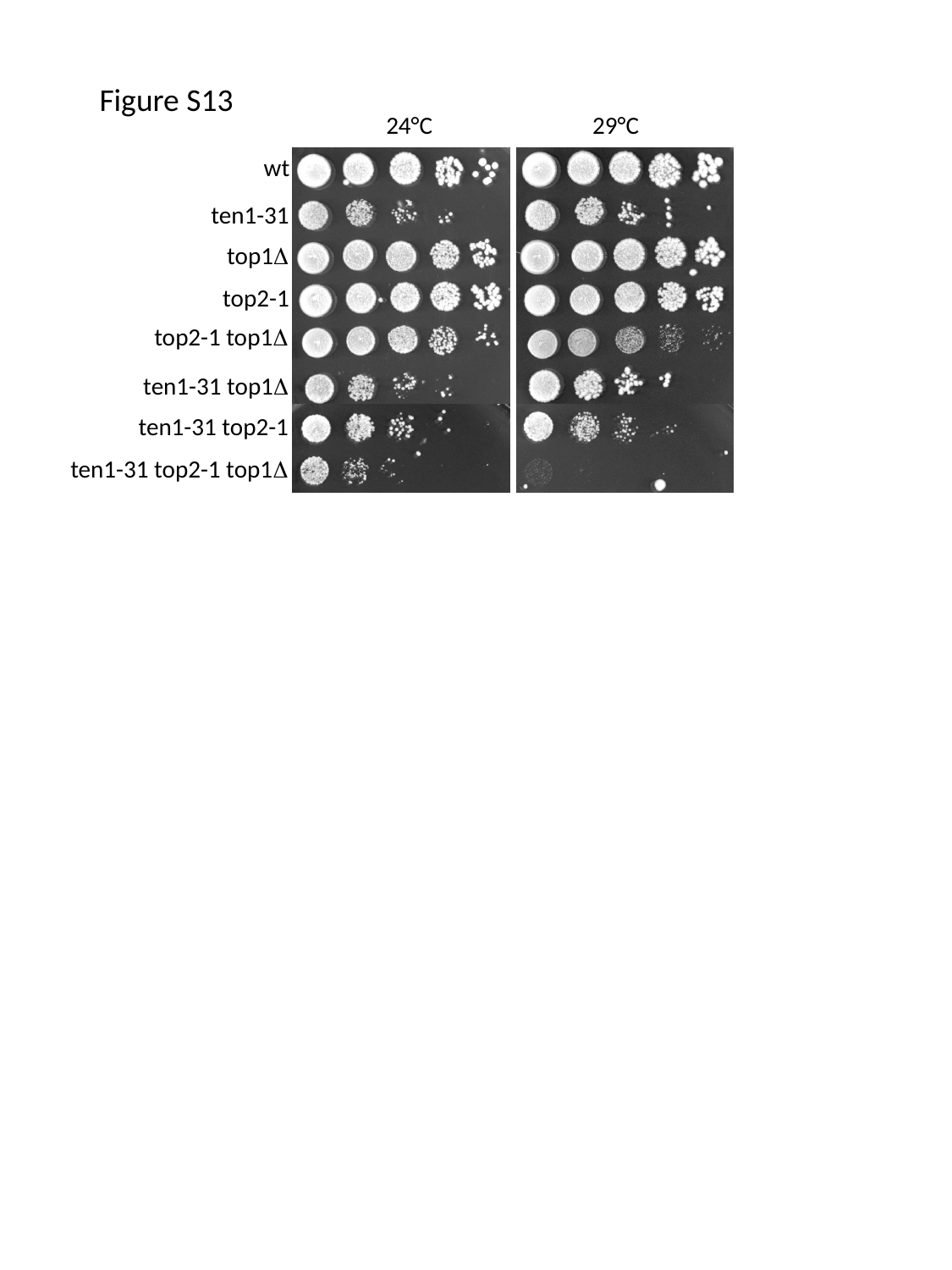

Figure S13
24°C
29°C
wt
ten1-31
top1Δ
top2-1
top2-1 top1Δ
ten1-31 top1Δ
ten1-31 top2-1
ten1-31 top2-1 top1Δ

### Supplementary file 15

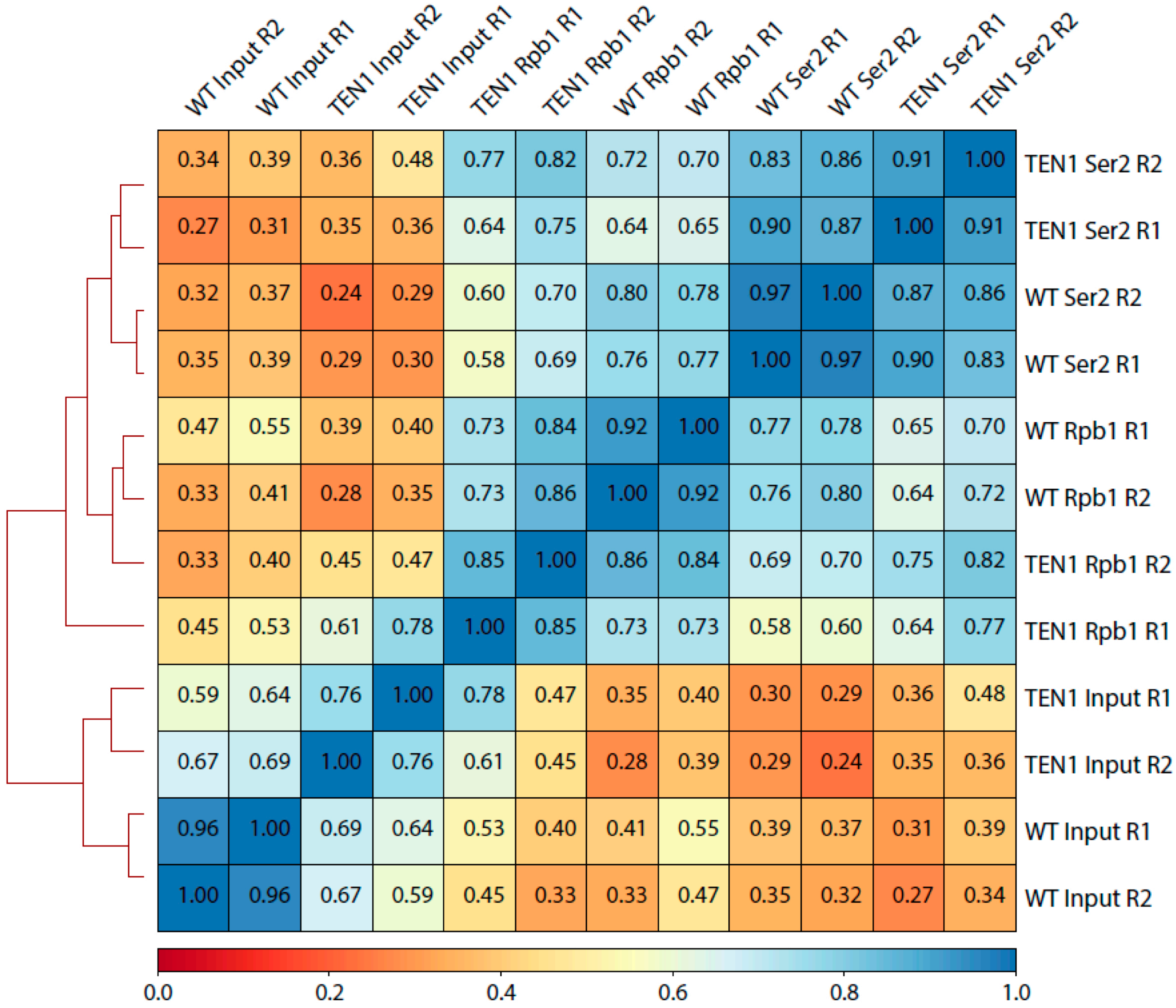
