## Supplementary material for "The telomeric Cdc13-Stn1-Ten1 complex regulates RNA polymerase II transcription"

### Slide 1
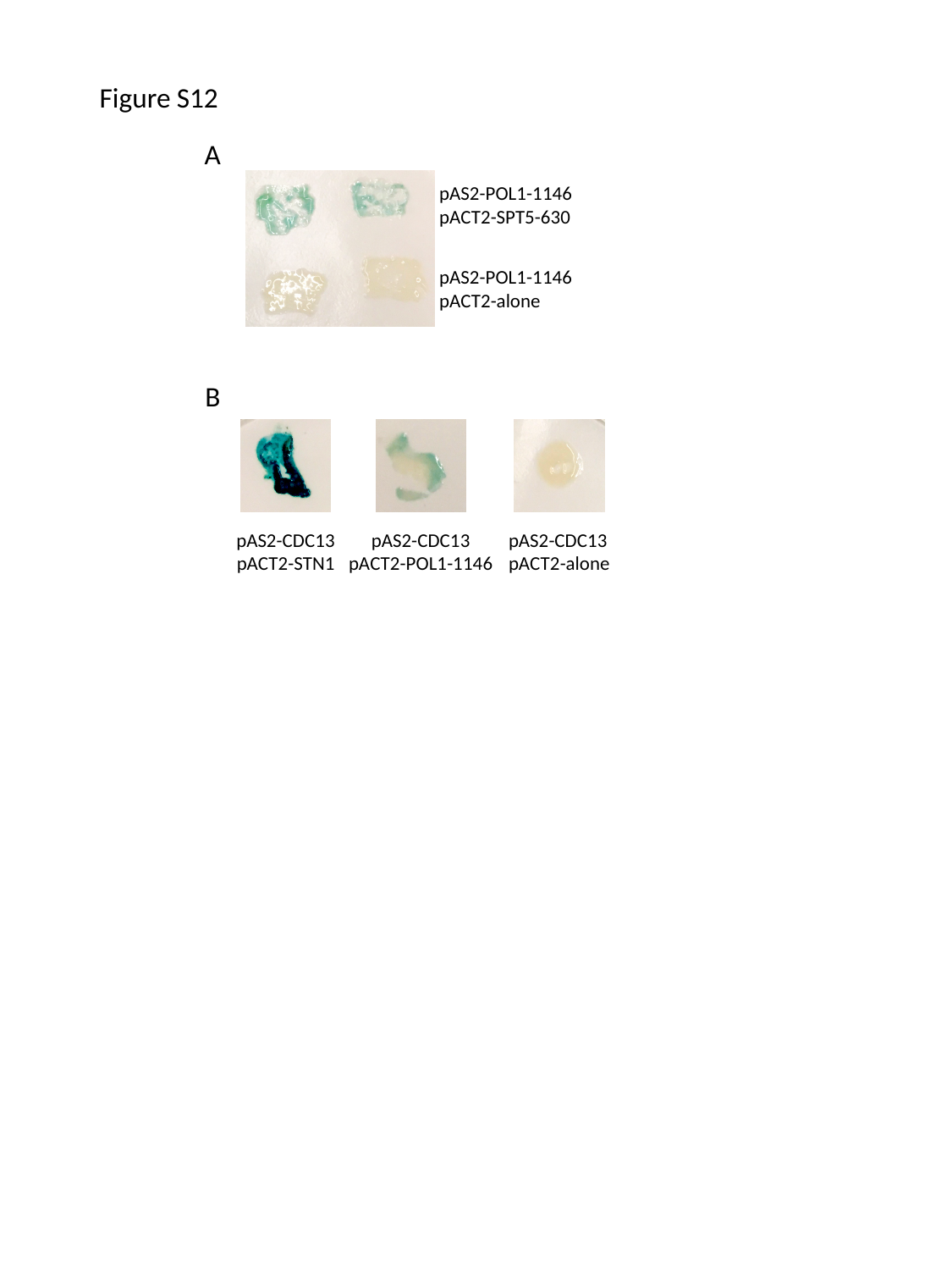

Figure S12
A
pAS2-POL1-1146
pACT2-SPT5-630
pAS2-POL1-1146
pACT2-alone
B
pAS2-CDC13
pACT2-STN1
pAS2-CDC13
pACT2-POL1-1146
pAS2-CDC13
pACT2-alone
