## Supplementary material for "The telomeric Cdc13-Stn1-Ten1 complex regulates RNA polymerase II transcription"

**Supplemental Table S1:** RNA-seq dataset info and alignment metrics

| **Sample** | **Reads** | **Trimmed reads** | **Percentage overall alignment rate** |
| --- | --- | --- | --- |
| TEN1 | 19166871 | 19123380 | 94 |
| TEN2 | 16053020 | 16014580 | 94.3 |
| TEN3 | 21265025 | 21216518 | 94.3 |
| WT1 | 21949183 | 21904287 | 95.5 |
| WT2 | 18590256 | 18538997 | 95.2 |
| WT3 | 21145339 | 21093191 | 95.1 |
| **Mean** | **19694949** | **19648492** | **94.7** |
