## Supplementary material for "The telomeric Cdc13-Stn1-Ten1 complex regulates RNA polymerase II transcription"

**Supplemental Table S2:** ChIP-seq dataset info and alignment metrics

| **Sample** | **Reads** | **Trimmed reads** | **Percentage overall alignment rate** |
| --- | --- | --- | --- |
| WR11 | 35606817 | 34285492 | 89.3 |
| WR12 | 24035727 | 22155522 | 62.87 |
| TR11 | 34037702 | 32789532 | 85.66 |
| TR12 | 22605480 | 21082236 | 83.62 |
| WS21 | 22216324 | 20630009 | 78.71 |
| WS22 | 20870427 | 19209542 | 80.02 |
| TS21 | 22365034 | 21660918 | 78.12 |
| TS22 | 21773279 | 20211331 | 81.87 |
| WIN1 | 32741704 | 31999784 | 96.13 |
| WIN2 | 24239678 | 22787207 | 96.22 |
| TIN1 | 30834426 | 30144680 | 96.72 |
| TIN2 | 23180827 | 20086273 | 97.38 |
| **Mean** | **26125618** | **24753543** | **85.55** |
